# Whole-body simulation of realistic fruit fly locomotion with deep reinforcement learning

**DOI:** 10.1101/2024.03.11.584515

**Authors:** Roman Vaxenburg, Igor Siwanowicz, Josh Merel, Alice A. Robie, Carmen Morrow, Guido Novati, Zinovia Stefanidi, Gert-Jan Both, Gwyneth M. Card, Michael B. Reiser, Matthew M. Botvinick, Kristin M. Branson, Yuval Tassa, Srinivas C. Turaga

## Abstract

The body of an animal influences how the nervous system produces behavior. Therefore, detailed modeling of the neural control of sensorimotor behavior requires a detailed model of the body. Here we contribute an anatomically-detailed biomechanical whole-body model of the fruit fly *Drosophila melanogaster* in the MuJoCo physics engine. Our model is general-purpose, enabling the simulation of diverse fly behaviors, both on land and in the air. We demonstrate the generality of our model by simulating realistic locomotion, both flight and walking. To support these behaviors, we have extended MuJoCo with phenomenological models of fluid forces and adhesion forces. Through data-driven end-to-end reinforcement learning, we demonstrate that these advances enable the training of neural network controllers capable of realistic locomotion along complex trajectories based on high-level steering control signals. We demonstrate the use of visual sensors and the re-use of a pre-trained general-purpose flight controller by training the model to perform visually guided flight tasks. Our project is an open-source platform for modeling neural control of sensorimotor behavior in an embodied context.

## Introduction

Animal behavior is the product of sensorimotor feedback control loops involving the animal’s brain, its body, and its environment [Merel et al., 2019b, Dickinson et al., 2000, Gomez-Marin and Ghazanfar, 2019, Chiel and Beer, 1997]. The body dictates precisely how motor commands from the nervous system are translated into action, and how the consequences of motor actions are reported via sensory feedback. Thus a detailed biomechanical understanding of how the body works is an important step towards understanding the neural control of movement. Here, we provide a new framework for physics simulation of an anatomically detailed biomechanical model of the fruit fly body. We validate the generality of our model by demonstrating realistic simulation of locomotion – both flight and walking – using reinforcement learning. Our general-purpose simulation framework is a platform for future brain-body modeling of diverse fruit fly behaviors.

We developed our fly body model in the open source MuJoCo [Todorov et al., 2012] physics engine. We used high-resolution confocal microscopy to image and construct a model of a female fruit fly (Fig. 1). To enable accurate modeling of fly behaviors such as flight and walking, we added new features to the MuJoCo physics engine. First, we developed a new computationally efficient, phenomenological fluid model to simulate forces resulting from fly wings flapping in air. Second, we developed adhesion actuators to model forces generated by insect feet gripping a surface. We validate the body model and the physics simulation by demonstrating realistic locomotion — both flight (Fig. 2) and walking (Fig. 3) — along complex natural trajectories with closed-loop sensorimotor control. In particular, by using end-to-end reinforcement learning (RL) to train a closed-loop sensorimotor controller to imitate the locomotor patterns of real flies, we validate the physics modeling by showing that we can replicate the body trajectories of real flies with similar leg and wing kinematics. We trained “steerable” low-level closed-loop sensorimotor controllers [Merel et al., 2019b] for walking and flight which can drive the fly body to generate complex trajectories using only high-level steering commands. And finally, we demonstrate the reuse of a pre-trained low-level flight controller by training the model fly to perform visually guided flight tasks (Fig. 4).

**Figure 1:**
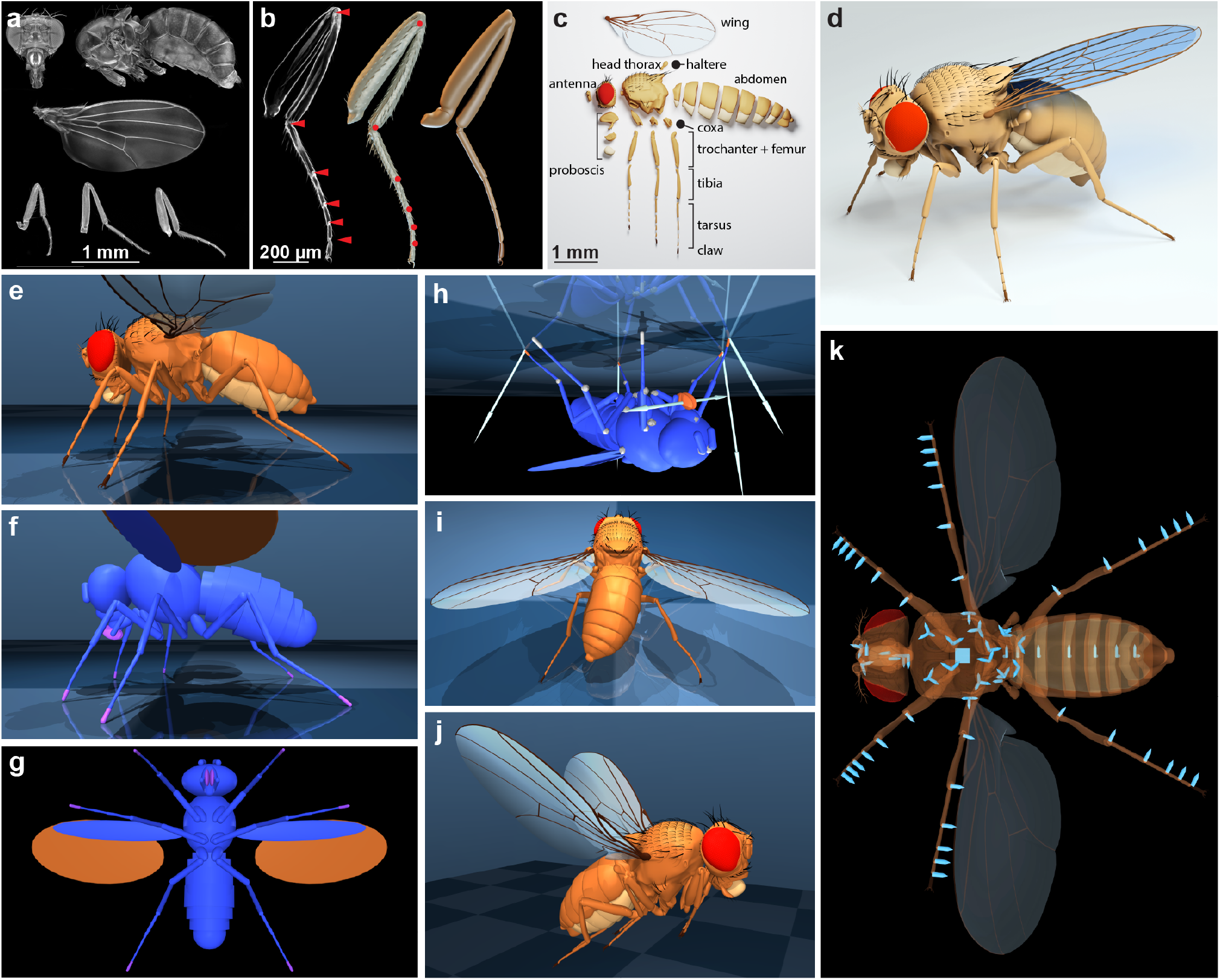
Constructing a 3D model of a female fruit fly from confocal data. (**a**) Compilation of 6 datasets representing a single fly; maximum intensity projections of confocal stacks showing head, thorax + abdomen, and legs. (**b**) A partial projection of the mid-leg confocal stack with the joints between femur-tibia and tarsal segments indicated by red triangles (left), a 3D mesh extracted from the stack (middle) and a low-polygon model of the leg (right). (**c**) An exploded simplified model (∼20k faces) showing body segments. (**d**) The complete anatomical model in the rest pose. (**e**) Side view of the body model in MuJoCo. (**f, g**) Side and bottom view of the geometric primitive (geom) approximation of body segments used to efficiently detect collisions and simulate physics. Blue: regular collision geoms. Mauve: Geoms that also have associated adhesion actuators. Ochre: Wing ellipsoids used by the advanced fluid model to enable flight. (**h**) Visualization of actuator forces generated when the model fly hangs upside down. The adhesion actuators of the front-right, middle-left, hind-right legs are actively gripping the ceiling (orange) and the labri (mouth) adhesors are also active; other actuators are inactive (white). The arrows, which visualize the contact forces, are proportional (and opposite) to the applied adhesion forces, minus gravity when applicable. (**i**) Exaggerated posture showing the result of activating the abdominal abduction and back-right tarsal flexion actuators. Abdominal joints and tarsal joints are each coupled with a single actuator (“tendon”) which simultaneously actuates multiple degrees of freedom (DoFs). (**j**) Model in a flight pose with legs retracted. (**k**) Bottom view, translucent visual geometry with light-blue arrows indicating joints. Cube: 6-DoF free joint (required for free center-of-mass motion in the simulator and is not a part of fly’s internal DoFs). Arrows: hinge joints; point in the direction of positive rotation. Groups of three hinge joints effectively form ball joints.

**Figure 2:**
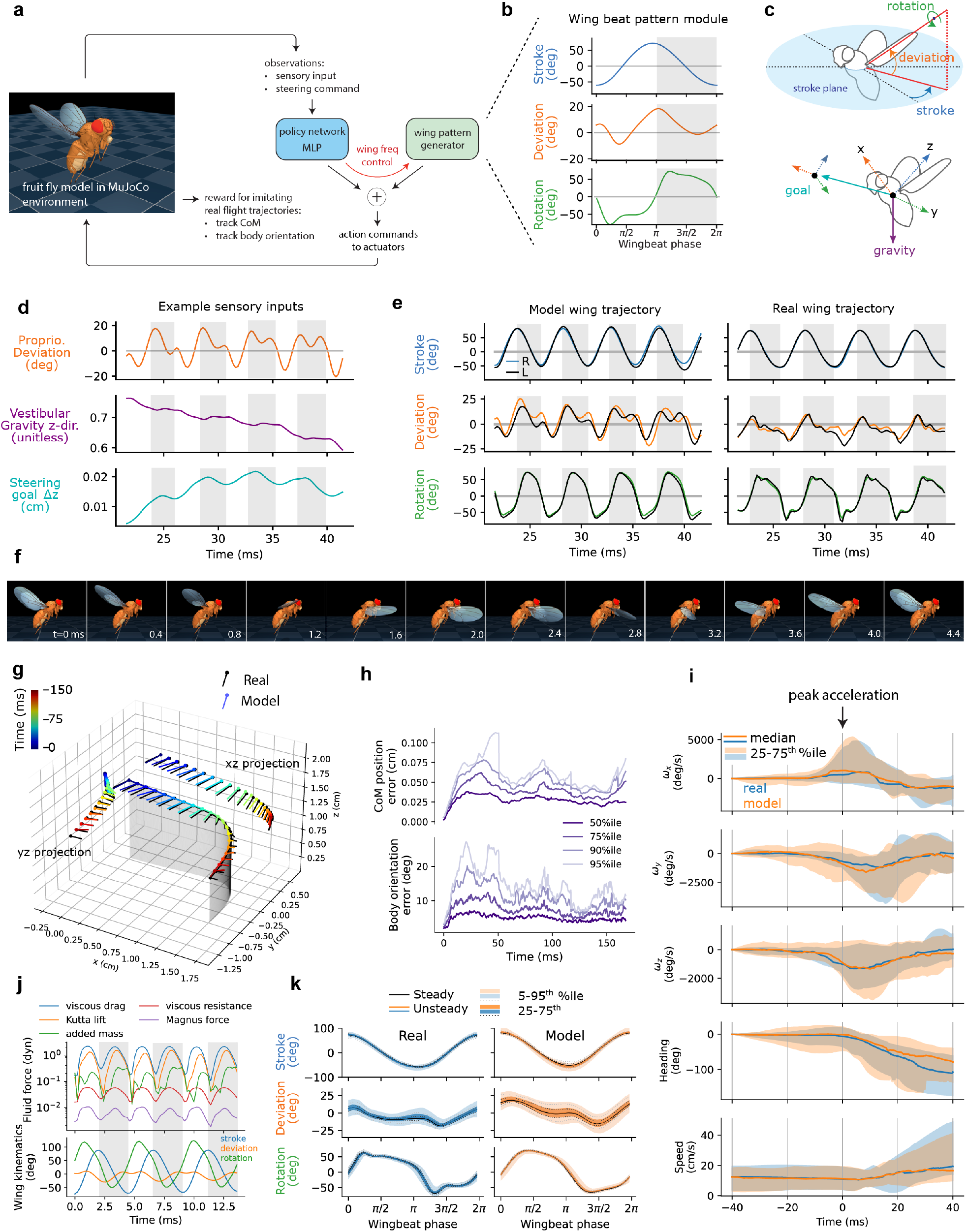
Flight imitation. (**a**) RL task setup. A single policy network is trained to imitate the body CoM and orientation across a data set of 216 trajectories (∼43 seconds total) of flying *Drosophila*. The controller consists of a trainable multi-layer perceptron (MLP) and a fixed wing-beat pattern generator (WPG). The net motor command received by the fly model is the MLP and WPG outputs summed. (**b**) One period of baseline wingbeat pattern (same for both wings) produced by the WPG. Gray stripe indicates wing downstroke. (**c**) Top: illustration of the wing coordinate system and wing angle names. Bottom: illustration of the body coordinate system and some of the sensory information available to the model: the direction to the goal CoM position and the direction of gravity. (**d**) A subset of policy inputs: wing deviation angle (proprioception), z-component of gravity direction (vestibular), z-component of target CoM position (steering command) for an example flight trajectory. (**e**) Wing angles during a saccade produced by the model and the corresponding real fly used for steering. Gray stripes indicate wing downstrokes. (**f**) Filmstrip of one full wing beat cycle of the model flying straight at 30 cm/s, shown with 0.4 ms step between frames. (**g**) Wings produce body movements through a phenomenological fluid model. Plot of the real fly’s body pose (black) and the model fly’s body pose traversing a test trajectory, colored by time. Projections of these on the xz and yz planes are also shown. Circles: head, lines: tail. (**h**) Percentiles of errors between the model and corresponding real fly’s body center-of-mass (top) and orientation (bottom) for 56 test trajectories. (**i**) Median and 25th-75th percentiles of body angular velocity, heading, speed for real and model flies during saccades in test set. The trajectories are aligned to peak acceleration at *t* = 0. Roll, *ω*_*x*_, and pitch, *ω*_*y*_, angular velocities are similarly important in model flies’ as real flies’ turns. A small divergence between the model and real flies occurs after the saccade. (**j**) Fluid model forces exerted on the left wing during a stable forward flight at 30 cm/s. The corresponding wing kinematics is shown at the bottom. (**k**) Percentiles of wing angles during steady (small body acceleration) and unsteady (large body acceleration) wing beats for model and real flies in the test set. Large body accelerations are achieved by similarly small alterations to the median wing beat pattern.

**Figure 3:**
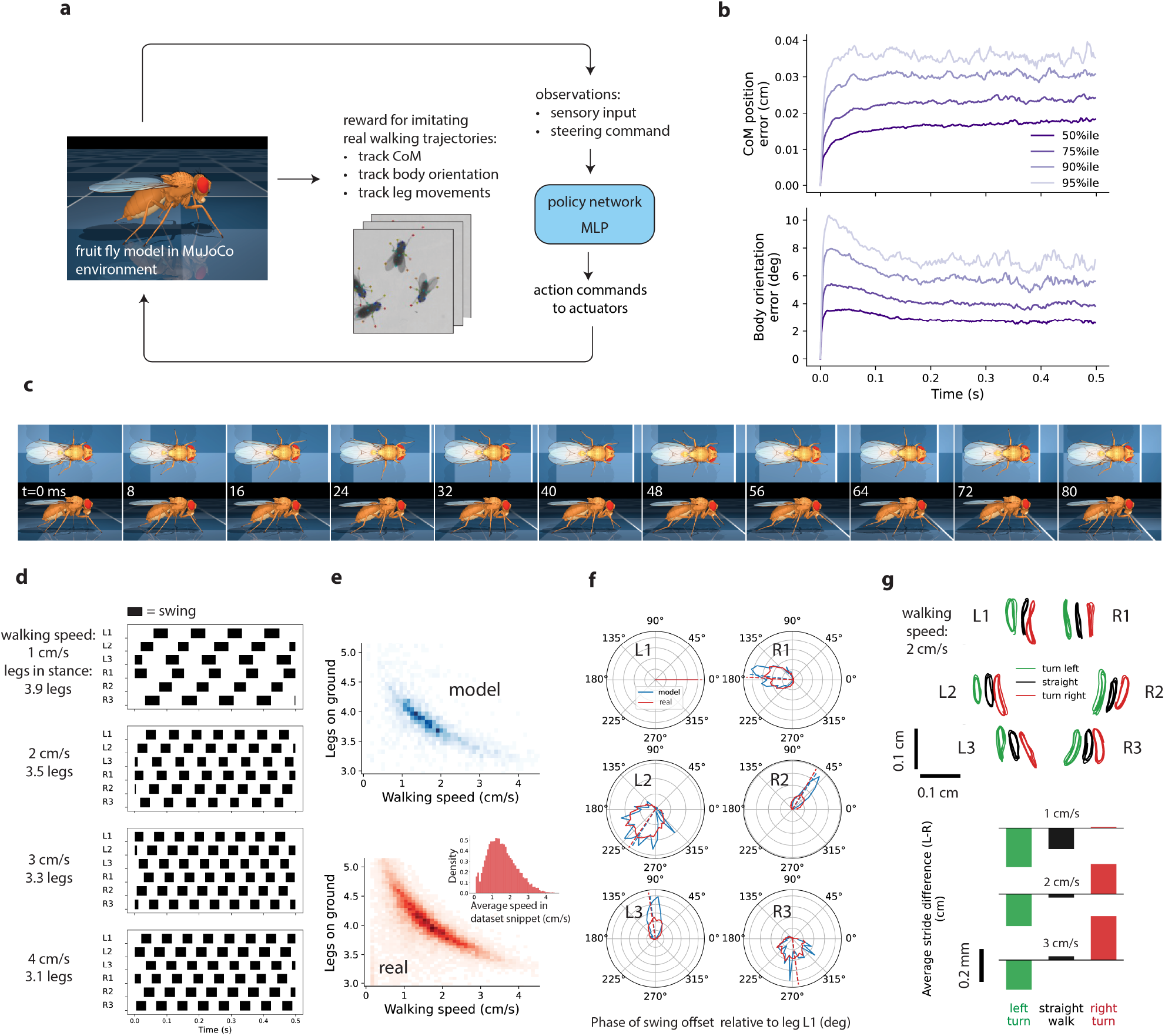
Walking imitation. (**a**) RL task setup. A single policy network is trained to imitate a dataset of 13k walking snippets (∼64 minutes total) of real *Drosophila*. Full body movements are imitated, including tracking body CoM position and orientation, and detailed leg movements. (**b**) Percentiles of errors between the model and corresponding real fly’s body center-of-mass (top) and orientation (bottom) for 3.2k test walking trajectories. (**c**) Filmstrip of one full leg cycle of the model walking straight at 2 cm/s with 8 ms step between frames. (**d**) Gait diagrams of fly model tracking artificial fixed-speed straight-walking trajectories at four speeds. For each speed, average number of legs simultaneously in stance position (on ground) is indicated. Black stripes indicate swing leg motion. (**e**) Number of legs simultaneously in stance position (on the ground) averaged over snippet vs average snippet walking speed. Top: model tracking all test set trajectories. Bottom: all of the walking dataset. Inset shows the distribution of average walking speeds per snippet in the dataset. (**f**) Distributions of phases of swing onsets of all legs relative to front left leg L1 in walking trajectories with mean speed in range [1.2, 1.7] cm/s. Blue is the model tracking testset trajectories and red is all of the reference data. Dashed lines indicate circular medians. (**g**) Learned turning strategy. Top: *xy*-projection of leg tip trajectories in egocentric frame for model walking straight, turning left, right, at speed of 2 cm/s. Leg tip trajectories are shifted horizontally for clarity. Bottom: Difference between left and right leg tip swing length averaged over all legs at different walking speeds.

**Figure 4:**
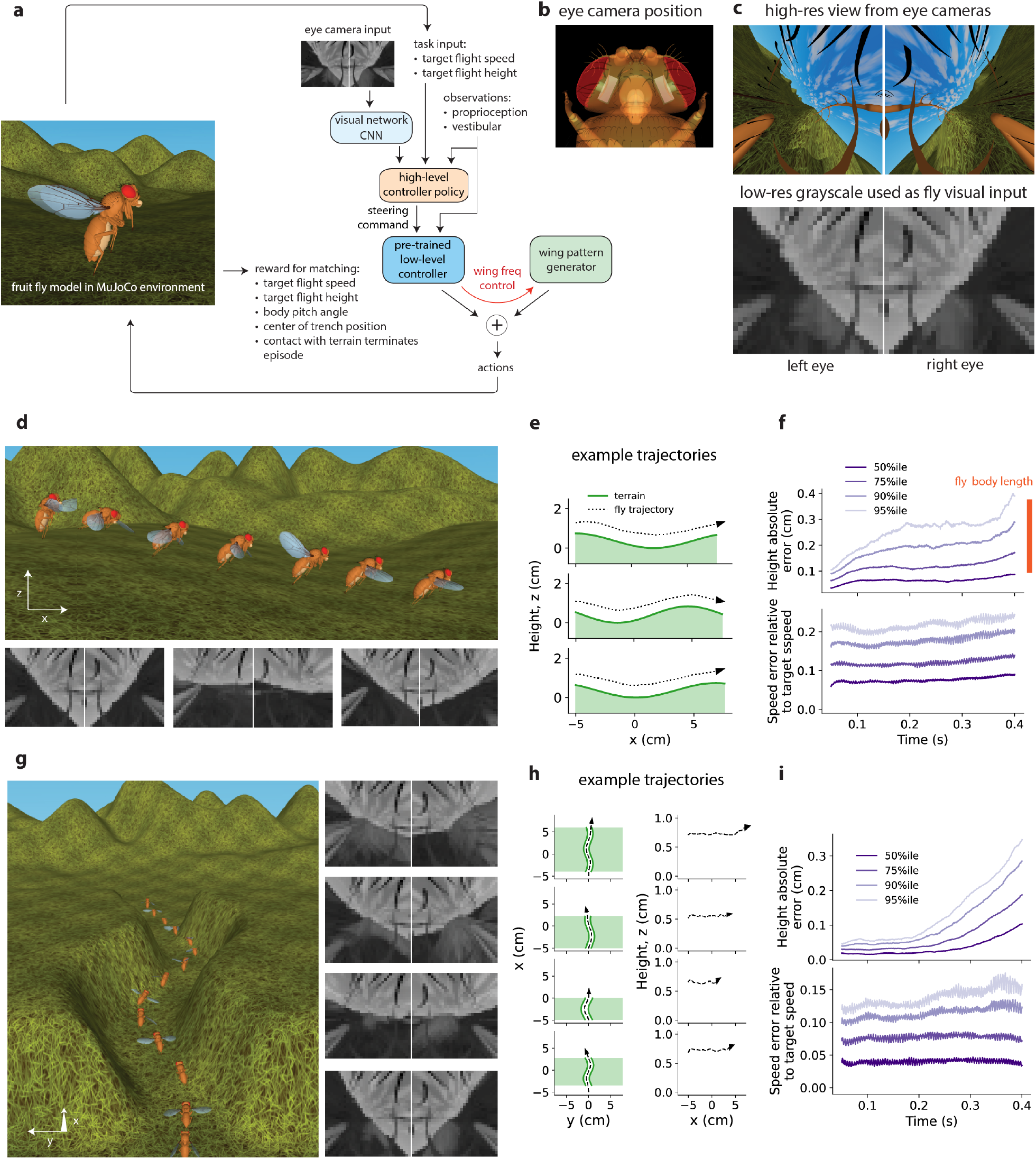
Vision-guided flight tasks: altitude control (“bumps”) and obstacle avoidance (“trench”). (**a**) RL task setup. Policy is trained to carry out forward flight at a given target speed and height while avoiding contact with terrain. In each training trajectory, the terrain is procedurally generated and the target speed and altitude are randomly selected. We considered two kinds of terrain. In the “bumps” task, the terrain is a sequence of sine-like bumps perpendicular to the flight direction. To solve the task of flying at given target height above the bumpy terrain, the agent has to learn to constantly adjust the flight altitude. In the “trench” task, the terrain is in the shape of a sine-like trench. To fly through the trench, the agent learns to execute turning maneuvers. To solve either task, the agent has to utilize the visual input from eye cameras to avoid terrain, and maintain target altitude. As in the flight imitation task, the flight controller consists of policy network and wingbeat pattern generator. The visual input is processed by an added convolutional network. The target flight speed and height are provided to the policy as task inputs. Both the policy and the convolutional network are trained end-to-end with RL. (**b**) Semi-transparent top-down view of fly model head showing schematically the position of MuJoCo eye cameras. (**c**) Fly visual input. Top: High-res view from eye cameras for fly’s flight position in Panel A. Bottom: Corresponding downsampled grayscale frames are used as fly visual input. (**d**) Top: Time-lapse of flight executed by policy trained in “bumps” task. Bottom: visual input frames captured by eye cameras for the first three time-lapse positions. (**e**) “Bumps” task performance: side-view of representative fly model trajectories. (**f**) Percentiles of height and speed errors for 1000 trajectories. The model body length (0.29 cm) is shown for comparison (**g**) Left: Time-lapse of flight generated by policy trained in “trench” task. Right: visual input frames for the first four time-lapse positions. (**h**) “Trench” task performance: top-down view of representative fly model trajectories (left) and their corresponding flight height (right). (**i**) Percentiles of height and speed errors for 1000 trajectories.

Our biomechanical model and physics simulation follows previous work on physics simulation of the worm [Boyle et al., 2012], hydra [Wang et al., 2023], rodent [Merel et al., 2019a] and fruit fly [Reiser et al., 2004, Dickson et al., 2008, Lobato-Rios et al., 2022, Wang-Chen et al., 2023, Melis et al., 2024] with qualitative improvements to the comprehensiveness and realism of the anatomy, biomechanics, physics, and behavior, leading to realistic simulation of both walking and flight in a single detailed general-purpose body model. The pioneering Grand Unified Fly [Dickson et al., 2008] demonstrated sensorimotor closed-loop visually guided flight behavior, with a reduced body model and a hand-designed controller. More recently, [Melis et al., 2024] modeled the wing hinge and its actuation in detail, revealing the physics of wing actuation by muscles. In parallel, NeuroMechFly [Lobato-Rios et al., 2022, Wang-Chen et al., 2023] introduced an anatomically detailed biomechanical body model capable of walking and grooming. A heuristically designed low-level walking controller was paired with a learned high-level controller to generate sensory-guided behaviors [Wang-Chen et al., 2023]. In this work, we developed a physics environment capable of both flight and walking with a single general-purpose, anatomically realistic body model. We used high-speed video based kinematic tracking [Branson et al., 2009, Pereira et al., 2022, Gosztolai et al., 2021, Karashchuk et al., 2021] to measure natural locomotion [Muijres et al., 2014, Muijres et al., 2015], and trained closed-loop low-level controllers capable of imitating arbitrarily complex natural trajectories using end-to-end reinforcement learning. As we demonstrate through inverse kinematics, our body model is capable of a rich repertoire of fly behaviors beyond locomotion, including grooming.

Our work lays the foundation for studying the neural basis of sensorimotor behaviors in the adult fruit fly. Our general-purpose and open-source model can be extended in the future to increase the realism of the biomechanics, neural control, and behavior as new data becomes available. For instance, new imaging of musculature and sensory systems can improve biomechanical fidelity of actuation and sensory feedback. The black-box artificial neural network controllers we trained using imitation learning can be made more realistic by introducing connectome constraints [Lappalainen et al., 2024] based on recent measurements of connectivity of the brain [Dorkenwald et al., 2023, Schlegel et al., 2023] and ventral nerve cord [Lesser et al., 2023, Azevedo et al., 2022, Cheong et al., 2023, Marin et al., 2023, Takemura et al., 2023], as well as neural activity [Turner et al., 2022, Pacheco et al., 2021, Aimon et al., 2023]. Thus our present work serves as a platform to enable future studies of embodied cognition [Zador et al., 2023], modeling the neural and biomechanical basis of sensorimotor behavior in unprecedented detail. Our fly model is available at https://github.com/TuragaLab/flybody.

## Results

### Body model geometry

We used confocal fluorescence microscopy to image the entire adult female fly body with high resolution (Fig. 1a, Methods, Supp. Data). Fluorescence microscopy enabled the staining of chitin to easily segment the shapes of body segments and identify the pivot points of all joints (Fig. 1b). Aberration-free high-resolution imaging of the entire body required disassembling it into smaller parts, chemical elimination of soft tissue, and bleaching of pigmentation (Methods). The dataset also enables the identification of origin and insertion sites of muscles, locations of the proprioceptive hair plates of the neck, coxae, trochanters, wing base and halteres — information that can be incorporated into future versions of the model.

We manually segmented 67 body components using Fiji [Schindelin et al., 2012], then simplified the corresponding meshes in Blender [Community, 2018], reducing the number of vertices by a factor of 1000, to allow efficient computational modeling while preserving morphological features relevant for our biomechanical study (Fig. 1b,c). In Blender, we assembled component meshes into a body by connecting them at 66 identified joint locations (Fig. 1b,d), amounting to 102 degrees of freedom (DoFs) determined in agreement with the literature (Fig. 1k). We modeled each biological joint as either a single hinge joint (1 DoF) or a combination of two or three hinge joints (2 and 3 DoFs, respectively). We estimated the joint angles corresponding to a resting fly pose (Fig. 1d,e) by visual inspection of videography. We note that these model joints are simplified approximations, particularly for the neck joint, wing hinge, and thorax-coxa articulation joint [Strausfeld et al., 1987, Melis et al., 2024, Gorko et al., 2024]. The fly model assembled in Blender defines the fly’s geometry and kinematic tree, and is available at https://github.com/TuragaLab/flybody.

### Modeling body physics

We exported the geometrical Blender fly model to MuJoCo and added the components required for physics simulation. This involved several steps (Supp. Section A). First, we generated primitive “geom” representations of each body part (Fig. 1f,g), which are used for efficient physics simulation and collision detection. Second, we measured the masses of each body part (Methods, Table 1). MuJoCo used these masses, along with an assumption of uniform density within each body part, to estimate the body part moments of inertia. Third, we added actuators to drive all the joints. Torque actuators were used for the wing joints, and position actuators for the rest of the body parts (Methods, Table 3). Since comprehensive measurements do not yet exist to faithfully model anatomically realistic muscle actuation, the choice of whether to use idealized position actuation or torque actuation is a matter of convenience and can be readily reconfigured.

**Table 1:** Empirical masses of the fly body parts. Averaged over 52 female flies.

| Body part | Mass (mg) |
| --- | --- |
| head | 0.15 |
| thorax | 0.34 |
| abdomen | 0.38 |
| leg (each) | 0.0162 |
| wing (each) | 0.008 |
| fly total | 0.983 |

Joint limits were determined by using inverse kinematics to match a diverse collection of poses from videography data, including grooming postures in which flies showcase their remarkable joint flexibility (Methods). Typically, each actuator applies force to a single DoF. However, in the multisegmented tarsi and abdomen body parts, several DoFs are coupled by a tendon and actuated by a single actuator to produce coordinated bending (Fig. 1i). Insect tarsi can adhere to surfaces, allowing flies to walk on walls and ceilings [Arzt et al., 2003]. To model this, we introduced adhesion actuators to MuJoCo. The adhesion actuators can simulate both active (controlled) and passive (uncontrolled) forces, and can now be used more generally for other models as a new MuJoCo feature (see Extended Data Fig. 7a for details). We added adhesion actuators to the tarsal tips (Fig. 1f,h). We also added adhesion actuators to the labrum to enable modeling of feeding and courtship behaviors [McKellar et al., 2020]. Fourth, we equipped the model with a sensory system, comprising vision, vestibular, proprioceptive, and mechanosensitive sensors, see Table 2 for details and Table 20 for correspondence between the sensory systems of our model and the real fly.

**Table 2:**
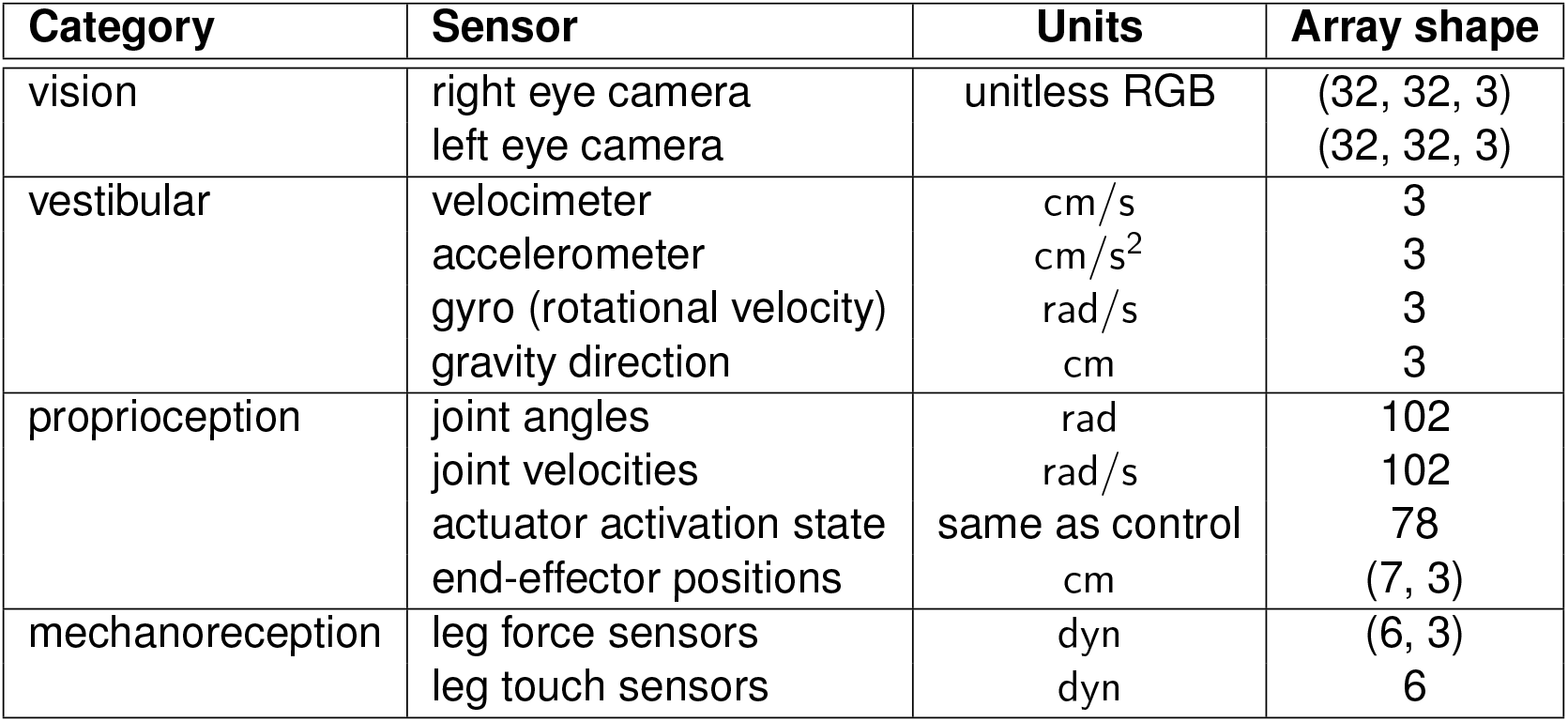
Sensory system of the fly model in its default configuration. Task-specific model modifications (e.g., disabling legs for flight) may include disabling some of the sensors and altering the dimensions of the proprioception observables. All observables are calculated in fly’s egocentric reference frame. The actuator activation state units are the same as the corresponding actuator control units.

| Category | Sensor | Units | Array shape |
| --- | --- | --- | --- |
| vision | right eye camera | unitless RGB | (32, 32, 3) |
|  | left eye camera |  | (32, 32, 3) |
| vestibular | velocimeter | cm/s | 3 |
|  | accelerometer | cm/s<sup>2</sup> | 3 |
|  | gyro (rotational velocity) | rad/s | 3 |
|  | gravity direction | cm | 3 |
| proprioception | joint angles | rad | 102 |
|  | joint velocities | rad/s | 102 |
|  | actuator activation state | same as control | 78 |
|  | end-effector positions | cm | (7, 3) |
| mechanoreception | leg force sensors | dyn | (6, 3) |
|  | leg touch sensors | dyn | 6 |

The result of these steps is an anatomically detailed, functional, biomechanical model of the entire fly body. All aspects of this model are programmatically modifiable. For instance, at simulation time, certain DoFs can be frozen (e.g., removing leg DoFs during flight), actuators can be programmatically enabled or disabled, body parts can be rescaled, etc. Similarly, sensors can also be enabled or disabled, and their properties (e.g., filtering in time) modified. The model is also extendable. For instance, the sensory system capabilities can be extended by adding biological realism with muscle actuation and detailed sensory transduction as more data becomes available.

### Modeling body-environment interactions

In addition to simulating the physics of the body itself, the MuJoCo physics engine also simulates the interactions of the body with its environment. These interactions include forces due to physical contacts, interactions with a surrounding fluid, e.g. air, as well as gravity. Contacts can be modeled both between different body parts of the fly, and between the fly body and its environment which is important for determining ground reaction forces during walking on a substrate.

Fluid interactions consist of forces generated by interactions of the wings with air. Accurate mechanistic simulation of fluid dynamics is a famously challenging computational problem. We therefore developed a phenomenological stateless quasi-steady state approximation of the fluid dynamics (Methods). Our fluid model is a generalization of [Andersen et al., 2005] to three dimensions. We model the forces and torques exerted on ellipsoid-shaped bodies moving through an incompressible quiescent fluid and approximated five fluid dynamics phenomena: added mass [Lamb, 1932, Tuckerman, 1925], viscous drag [Duan et al., 2015], viscous resistance [Stokes, 1850], Magnus lift [Seifert, 2012], and Kutta lift [Kutta, 1902]. The forces and torques arising from rotation and translation of an ellipsoid body are approximated by polynomial functions of fluid parameters (density, viscosity) and ellipsoid parameters (shape, size, mass, linear and angular velocities), with a slender ellipsoid approximation of the wings. This fluid model is another new MuJoCo feature developed to facilitate our study, and can be used more generally for other models.

In order to accurately generate flight with the body model, our phenomenological model must to account for the total force resulting from the movement of real wings through air regardless of mechanism, including e.g. passive contributions from wing flexibility [Sane, 2003]. Therefore in order to compensate for all the ways in which in our approximation deviates from the true forces, we optimized the coefficients for the magnitudes of each of the fluid components until stable hovering was achieved (Methods). In our fluid model for the fly, we found that Kutta lift and viscous drag components dominated (see Flight imitation section below). Kutta lift is the lift force generated by circulation of the fluid around the wing, while viscous drag is a force opposing movement of the wing through the fluid. In practice, we found that the coefficients for these two components did not require fine-tuning and similar flight performance resulted from coefficients varied by up to 20% (Extended Data Fig. 5). We should note that our quasi-steady state approximation ignores the dynamical effects of turbulence, vortices and other stateful-fluid phenomena which are known to contribute to insect flight [Sane, 2003], while only capturing their time-averaged contributions.

Simulation of behavior by the numerical integration of the passive and active dynamics of the fly model and its interactions with the environment is a considerable computational challenge, both in terms of the dimensionality of the system and the small integration time step required for numerically accurate simulation. Our fly body model is of considerable complexity (102 DoFs) and the simulation must model fast behaviors, such as ∼ 200 Hz wing flapping during flight, requiring a small simulation time step, e.g. ∼ 0.1 ms. On a single core of an Intel Xeon CPU E5-2697 v3, the simulation of 10 ms of full flight and walking environment loops (Fig. 2a, Fig. 3a) took 421.5 ms and 68.65 ms, respectively, which is fast enough to enable reinforcement learning of motor control. The fraction of time used by the MuJoCo simulator alone (i.e., excluding the rest of the environment loop components: policy network and python RL environment overheads) was 55.5 ms and 23.2 ms for flight and walking, respectively (Supp. Section C, Table 21).

### Imitation learning of locomotion

We used deep reinforcement learning to train our model fruit fly to generate specific behaviors. In particular, we trained an artificial neural network to serve the role of the nervous system, forming a sensorimotor controller driving locomotion in a closed loop. At each time step, MuJoCo simulates signals from the sensory system which are provided as input to the neural network. The neural network then computes control signals corresponding to each actuator in the body, which are then used by MuJoCo to simulate the active forces generated by the body for that time step.

We used imitation learning [Peng et al., 2018, Merel et al., 2020a, Hasenclever et al., 2020], a data-driven approach to train our neural-network controllers, to generate realistic locomotion. Here, the network was trained so that the trajectory of the center of mass (CoM) position and orientation of the body, as well as individual body parts, matched the corresponding trajectory of a real fly during free locomotion measured from video. To train networks which can generalize from training trajectories to locomotion along novel trajectories, we trained “steerable” low-level controllers [Merel et al., 2019b]. These neural networks are analogous to, but not meant to exactly model, the ventral nerve cord (VNC) of the fly central nervous system and are responsible for translating high-level (descending) command signals from the central brain into low-level motor control signals. The high-level commands in our model correspond to the desired change in the fly’s (CoM) position and orientation (6 DoFs) at each time step, while the low-level motor control signals drive individual actuators in the body.

We trained two steerable neural network controllers (policy networks): one for flight (Fig. 2) and one for walking (Fig. 3). From training data in the form of real trajectories of the CoM and the body parts of walking and flying fruit flies (Methods), we used the CoM trajectory of the real fly as the high-level steering command to the network, and then constructed a reward function which rewarded the fly for matching both the target CoM as well as the locations of each body part at each time step. The reward is then maximized when the model fly tracks the CoM of the real fly, while moving its body parts in the same manner as the real fly. This form of imitation learning teaches the model fly to move its body along the same path as a real fly, using the same pattern of limb or wing movements as the real fly.

The steerable controller neural networks were feedforward multilayer perceptrons (MLP), which received egocentric vestibular and proprioceptive signals as input, in addition to the high-level steering commands. They were trained to generate motor control signals actuating the body in order to maximize the imitation reward using the DMPO agent, an off-policy model-free RL algorithm [Abdolmaleki et al., 2018b, Abdolmaleki et al., 2018a] with a distributional critic value function [Bellemare et al., 2017]. This is an actor-critic style algorithm which trains two networks: a policy network which learns a nonlinear mapping from the sensory observations to the control signals, and a critic network which predicts the expected discounted cumulative reward resulting from applying a given control signal at a given state. We used the implementation of DMPO from the acme [Hoffman et al., 2020] reinforcement learning python library. Training until convergence can require the order of ∼ 10^9^ (walking) and ∼ 10^8^ (flight) simulation steps with ∼ 10^8^ and ∼ 10^7^ of policy network weight updates, respectively. To reduce the training time to a practical range of days or hours, we developed a multi-CPU and GPU parallelization scheme [Horgan et al., 2018] using Ray [Moritz et al., 2017], a general-purpose distributed computing package (Methods). Our code is general purpose and open source. Other RL algorithms can be easily paired with our model to train controllers in a modular fashion. We describe the specifics of the two controllers next.

### Flight

We trained a steerable flight controller (Fig. 2a) with imitation learning using previously collected high-speed videography of freely flying *Drosophila hydei* performing spontaneous saccades [Muijres et al., 2015] and forced evasion maneuvers [Muijres et al., 2014]. These datasets combined contained 272 individual trajectories (∼53 seconds of flight) of the CoM along with wing kinematics during turns, speed and altitude changes, flying straight, sideways and backward, and hovering in place (Methods). While *Drosophila hydei* is a somewhat larger fly compared to *Drosophila melanogaster*, we expect that the body and wing kinematics are similar [Dickinson and Muijres, 2016], and nevertheless, these are the best available data. A single controller network was trained to imitate all 216 flight trajectories across the (train) dataset. This steerable controller could maintain stable flight and drive the fly to traverse new flight trajectories (Fig. 2).

The design of our flight controller policy network was inspired by the finding that flies use only subtle deviations from a nominal wing beat pattern to control their flight trajectories [Muijres et al., 2014, Muijres et al., 2015]. Our flight controller consisted of two parts: a fixed wing beat pattern generator (WPG) and a trainable fully-connected MLP (Fig. 2a). The WPG produce a fixed mirror-symmetric periodic baseline wing beat pattern (Methods), based on wing trajectories recorded from a hovering *Drosophila melanogaster* [Fry et al., 2005, Dickson et al., 2008] (Fig. 2b). The policy neural network controls the base frequency of the WPG as well as small wing motion deviations from the WPG baseline pattern, resulting in the full repertoire of flight behaviors from the training datasets. As the WPG’s baseline pattern was already close to the required wing motion solution, in effect the WPG also provided a good initialization which sped up training significantly.

In addition to the steering command, the policy network controller received a 62-dimensional sensory signal from the proprioceptive and vestibular sensory systems (Fig. 2d, Methods, Table 8). The controller output 12-dimensional instantaneous wing torques, head and abdomen angle change, and WPG frequency (Methods, Table 9). While not strictly necessary as this could be learned by imitation, the legs were retracted to their typical flight position (Fig. 1j) and their DoFs were frozen to reduce control complexity. The policy network was trained so that the model’s body CoM and orientation matched those of the reference trajectories. Note that reference trajectory wing angles were not used to train the model, but were only used for evaluation. The reward function and training details are provided in Methods.

We evaluated the performance of our trained steerable controller on a test set of 56 CoM trajectories. The model fly was able to match the desired CoM trajectory with high accuracy (median CoM position error of 0.25 mm, median body orientation error <5 °, Fig. 2h, example trajectory shown in Fig. 2g). A filmstrip of a single wing beat cycle produced by the trained model fly when commanded to fly straight at 30 cm/s is shown in Fig. 2f. The model fly was trained to match the target CoM trajectories of real flies, but was largely free to achieve this via wing trajectories only constrained (through the DMPO action penalization, Methods) to stay close to a common baseline WPG pattern. Therefore, we compared model and real wing trajectory patterns as a means to characterize the fidelity of our physics simulation of fluid forces as well as the realism of model fly behavior. We found that a good match in CoM trajectory was achieved by qualitatively similar wing trajectories of the model and real flies, but with slight differences in wing beat frequency (Fig. 2e). In general, we found that while the wing beat frequency changed during the maneuvers by up to ∼ 40 Hz in the datasets, in the model fly these frequency changes clustered more tightly in the ∼ 0 − 10 Hz range. As the model and data flies were different species with average wingbeat frequencies of 218 Hz (*Drosophila melanogaster*) and 192 Hz (*Drosophila hydei*), we did not attempt to quantitatively compare wing beat trajectories.

Similar to real flies, the model fly only generated small differences between the left and right wing trajectories in order to generate large accelerations during saccade turns (Fig. 2e). Since the model fly was trained and evaluated on trajectories involving such extreme maneuvers, we were able to compare changes in wing beat patterns between “steady” flight during periods of minimal acceleration, and “unsteady” flight during periods of large acceleration (Fig. 2k). Our model fly wing trajectories recapitulated previous observations [Fry et al., 2005, Dickson et al., 2008, Muijres et al., 2014, Muijres et al., 2015], that only small differences in the wing stroke between steady and unsteady flight (Fig. 2k) are sufficient to generate large changes in CoM trajectory (Fig. 2i). Further, the match in median angular velocities, heading, and speed, show that our model fly recapitulates many of the stereotypic features of the maneuvers used by real flies to turn [Muijres et al., 2015, Dickinson and Muijres, 2016].

Finally, we analyzed how forces were generated by our phenomenological fluid model to support flight (Fig. 2j). We found that just two components, the viscous drag and Kutta lift, dominated the force generation during a wing beat cycle. All other components were 1–2 orders of magnitude smaller.

### Walking

We trained a steerable closed loop controller for walking (Fig. 3a), also using imitation learning. We performed high-speed top-down-view videography (150 fps) of groups of fruit flies walking and interacting freely in a circular arena. Automated pose tracking was applied to female fruit flies to track the 2D locations of 13 keypoints located at the head, thorax, abdomen and 6 leg tips (Methods). The 3D pose of all the body DoFs cannot be unambiguously inferred from just these 2D keypoints locations, so we used a form of regularized inverse kinematics to infer an approximation of the full 3D fly pose trajectories for all degrees of freedom (Methods). The collected trajectories (∼16k trajectories, 80 minutes total) consisted of flies walking freely at variable speeds (∼0–4 cm/s, Fig. 3e inset), turning, and standing still.

Flies use a variety of gait patterns as a function of walking speed, characterized by differences in limb coordination and the number of feet on the ground at any given time [DeAngelis et al., 2019]. This is in contrast to flight where the full range of flying behaviors is executed by subtle modifications to a common baseline wingbeat pattern [Muijres et al., 2014, Dickinson and Muijres, 2016]. Because of the significant gait variability, we could not utilize a simple pattern generator for walking imitation as we did for flight. Instead, the controller consisted of a single fully-connected MLP (Fig. 3a) and was trained without enforcing any particular structure such as pattern generators. A single MLP policy network was trained on all ∼ 13k train set walking trajectories.

A much larger number of DoFs (59 for walking vs 12 for flight) has to be controlled during walking compared to flight, corresponding to the legs (including adhesion), abdomen, and head. Therefore, the sensory signals provided to the controller network were much higher (286-dimensional (Methods, Table 16)), corresponding largely to an increase in proprioceptive signals. Similarly, a 59-dimensional motor control signal was required from the controller (Methods, Table 17). While not strictly necessary as this could be learned by imitation, the wings were folded and their actuation disabled. During training, the model fly was rewarded for reproducing the detailed leg movements of the real fly, as well as tracking the CoM trajectory, in response to high level steering commands in the form of desired future body CoM position and orientation. As we only had measurements of the body kinematics and no direct reference measurements of leg adhesion, the reward function did not include a term for imitating leg adhesion. However, we observed that the agent generally learned to activate adhesion when legs were in stance (on the ground). See Methods for walking imitation reward and training details.

We characterized the walking behavior produced by the steerable controller on a test set of 3.2k trajectories. The CoM position tracked the desired position and orientation well (median position error: 0.4 cm, median orientation error: 4°, Fig. 3b). A single walking cycle produced by the model fly, commanded to walk straight at 2 cm/s, is shown in Fig. 3c. We then analyzed the gaits produced by the model fly when commanded to walk straight at a given constant speed. The gait diagrams (Fig. 3d) show the duration that each leg spent in stance or swing phases, defined by whether the leg was stationary on the ground or moving, respectively (see the distribution of walking speeds in data in Fig. 3e, inset). At any time, fruit flies generally have at least three legs on the ground in stance phase, but more legs can be in stance at slower walking speeds [DeAngelis et al., 2019]. The model fly recapitulates this phenomenon with 3.1 legs in stance phase on average at during high speed walking (4 cm/s), and 3.9 at the slower walking speed of 1 cm/s. Across the full range of walking speeds observed in the dataset, we find good agreement for average number of legs on the ground between model and real flies (Fig. 3e). We analyzed the leg coordination patterns of the model and real flies by computing the phase delay during a walking cycle between the swing phase onsets of each leg relative to the left foreleg (L1), and find good agreement between the phase distributions of the model and real fly legs (Fig. 3f).

Fig. 3g shows the learned turning strategy. We analyzed leg tip trajectories produced by the model fly when commanded to either walk straight (black) or turn left (green) or right (red) at a constant walking speed of 1, 2, or 3 cm/s and a fixed turning radius of 1 cm. The model fly decreases stride length of legs on the turning side and increases stride length of legs on the opposite side as seen in the leg tip projections for turning at 2 cm/s (top), and qualified for all speeds (bottom). This agrees with the observed change in step length for the middle- and hind-limbs of freely walking flies [DeAngelis et al., 2019]. In contrast, our model has learned an asymmetric pattern for the forelimbs, modulating the length for the front limbs during left turns but not right turns, a pattern not typically seen in walking measurements of real flies.

The addition of adhesion actuators enables simulation of complex 3D navigation on steep surfaces. To demonstrate this, we designed an imitation learning walking task requiring the fly model to move across hilly terrain with gradually varying steepness. We made sure that this task would be impossible to solve with simple gravity-based leg-floor friction (Methods, Extended Data Fig. 7). We analyzed the adhesion actuation generated by the model and resulting adhesion and contact forces as a function of the angle of the surface. Within MuJoCo’s Coulomb friction model, adhesion forces are normal to the surface, pushing the fly legs towards the surface and resisting slip across the surface by effectively increasing the threshold tangential force at which slipping would occur. The model learns to vary the adhesion actuator force to provide sufficient slip-resisting forces depending on the terrain steepness. The model generated larger adhesion forces with the fore- and mid-legs on the upward slope and uses the hind-legs on the downward slope to resist slipping. Further details in Methods and Extended Data Fig. 7.

### Vision-guided flight with flight controller reuse

The fruit fly is a highly visual insect, with two compound eyes dominating its head and with two optic lobes which each comprise about a third of the fly’s brain.Therefore, in addition to proprioceptive and vestibular sensors, we also modeled visual sensors. We model the eyes with MuJoCo camera sensors (Fig. 4b), rendering a square lattice of 32×32 pixels with a field of view of 150 degrees. This roughly corresponds to the resolution of the *Drosophila* compound eyes with an inter-ommatidial (“pixel-to-pixel”) visual angle of 4.6 degrees [Zhao et al., 2022] (Fig. 4c). We demonstrate these sensors by training the model fly to perform two tasks (“bumps” and “trench”) which both require using vision to navigate. Fig. 4c shows an example of a low-resolution visual input frame and its high-resolution counterpart rendered during flight.

We reused the general-purpose steerable low-level flight policy trained in the flight imitation task above as a part of the vision-guided flight controller architecture, which we trained using end-to-end reinforcement learning (Fig. 4a). The flight controller consists of the pre-trained policy (and the WPG) from the flight imitation acting as a low-level controller directly controlling the wing motion and a high-level controller “navigator” policy, which only issues low-dimensional steering commands to the low-level controller. In addition to the 62-dimensional proprioceptive and vestibular sensory signal, the high-level controller network also received a low-dimensional visual feature representation produced by a convolutional network (CNN) applied to the visual inputs. The CNN module received a 6,144-dimensional visual signal from the eyes. The high-level controller also received an additional 2-dimensional task-specific input: target flight height and speed (Methods, Table 12). The low-level controller received the same 62-dimensional sensory input (but not the task input and not the visual input), as well as the steering command from the high-level controller. As in the flight imitation task, the low-level flight controller produces 12-dimensional wing torques, head and abdomen angles, and WPG frequency (Methods, Table 13). We kept the weights of the low-level controller network frozen and jointly trained the CNN representing the visual system and the high-level MLP producing the steering commands in an end-to-end fashion to maximize task reward.

In both tasks, the precise terrain of the virtual world, as well as the target height and speed were randomly generated in each training and test episode (Methods, Table 14). Contact with terrain results in early episode termination with failure (Methods). The model fly starts at zero speed, requiring that the fly speed up to the target speed at the beginning of the task.

### Bumps task

During flight, fruit flies use visual cues to estimate their altitude above the ground, which could be used to maintain a constant height above the ground when flying over uneven terrain [Straw et al., 2010]. We modeled this visually guided altitude control task by constructing a virtual world with a randomly generated sinusoidal height profile of the terrain. The model fly is rewarded for flying straight at a constant target forward velocity while maintaining a constant target height above the ground (Fig. 4d). After training, the model fly successfully learned to use its visual system to match the target altitude (Fig. 4e,f, median errors after the initial speed-up period: height error: 0.045 cm, speed error: 2.2 cm/s).

### Trench task

In a second task, we trained the fly to navigate through a trench without bumping into the walls of the trench. A virtual world was constructed with a trench with a sinusoidal curving profile (Fig. 4g) and constant width and depth. The model fly was rewarded for maintaining a constant forward speed and constant height inside the trench, while a collision with the trench walls resulted in an early demise and termination of the episode and a loss of future rewards.

Thus maximizing the reward requires that the model fly successfully navigate the trench by turning to avoid collisions with the walls. After training, the model fly successfully learned to use its visual system to detect the walls of the trench and navigate the entirety of the trench while maintaining constant target height and target speed (Fig. 4h,i, median errors after the initial speed-up period: height error: 0.032 cm, median speed error: -0.16 cm/s).

## Discussion

The behavior of an animal emerges from the interplay between its nervous system, its body, and its environment. Here, we demonstrated realistic complex locomotion behavior, both walking and flight, in an anatomically detailed whole body model of the fruit fly. This advance was enabled by improved simulation of the physics of the body and its interactions with the world, and by using deep reinforcement learning to approximate the nervous system with an artificial neural network trained to mimic the behavior of real flies. We constructed an anatomically detailed model of the fruit fly with 67 rigid body parts and 102 degrees of freedom, actuated via torques at the joints. Using the MuJoCo physics engine, we simulated the body and its interactions in its environment through rigid body collisions and through fluid interactions with air. Finally, we have used deep RL and imitation learning approaches to construct a closed-loop low-level neural controller that induces the body to generate realistic patterns of body movement, both for walking and flight modes of locomotion, along arbitrary trajectories. All three components are released as open source software along with pre-trained controllers, and can be accessed at https://github.com/TuragaLab/flybody/.

The present work integrates measurements across multiple spatial and temporal scales, bringing together microscopy of static anatomy with high speed videography of dynamic locomotor behavior. Our model accurately simulates what forces are generated by the body and what quantities are sensed by the body, through idealized actuators and sensors. We intend our open source project to serve as a platform which can be built upon in several ways. Confocal imaging (Extended Data Fig. 1), as well as micro CT [Lobato-Rios et al., 2022] and XNH [Kuan et al., 2020] imaging of the body support detailed musculo-skeletal measurements across the whole body which can be used to model anatomically detailed muscle actuation. This includes modeling of the neck [Gorko et al., 2024], wing hinge [Melis et al., 2024], and coxa [Kuan et al., 2020, Mamiya et al., 2023] joints. On the sensory side, the idealized sensory systems in our model can be improved by incorporating recent mapping of proprioceptive organs in the leg and wing [Kuan et al., 2020, Mamiya et al., 2023], and the recently developed eye-maps can be used to inform the precise positions of individual ommatidia in visual space [Zhao et al., 2022]. Further, model-based pose-tracking algorithms could be used to extract improved kinematic detail from existing high-speed single- and multi-view videography [Bolaños et al., 2021, Jiang and Ostadabbas, 2023, Plum et al., 2023, Sun et al., 2023].

Future work can leverage the connectomic mapping of the entire fruit fly nervous system [Dorkenwald et al., 2023, Schlegel et al., 2023, Lesser et al., 2023, Azevedo et al., 2022, Cheong et al., 2023, Marin et al., 2023, Takemura et al., 2023] to more realistically model neural circuits underlying sensorimotor behavior. Our body model enables the prediction of sensory inputs to and motor outputs from the nervous system during behavior on a moment-by-moment basis. These predictions can be combined with newly revealed connectomic understanding of the precise mapping between individual sensory and motor neurons and the body at the resolution of individual degrees of freedom [Azevedo et al., 2024]. Recent work by [Lappalainen et al., 2024] showed that connectome-constrained network models can be combined with a characterization of the input-output function of the network to enable predictions of neural activity at single neuron resolution across the network. Therefore, together with imitation learning, our model can be combined with measurements of the connectome and behavior to enable the model-based investigation of the neural underpinnings of sensory-motor behaviors such as escape invoked by looming stimuli [Card, 2012], gaze-stabilization [Cruz and Chiappe, 2023], and the control of locomotion by the ventral nerve cord. In the long term, our whole body model, in concert with a connectome of the whole nervous system and comprehensive measurements of behavior, as well as connectome-constrained deep mechanistic neural network modeling [Lappalainen et al., 2024, Mi et al., 2022], could eventually be used to construct whole animal models of both the entire nervous system and body of the adult fruit fly.

## Data availability

Confocal imaging stack, flight and walking imitation datasets, base wingbeat pattern, and trained controller networks are available at https://doi.org/10.25378/janelia.25309105

## Code availability

Fly model and code are publicly available at https://github.com/TuragaLab/flybody

## Methods

### Sample preparation for anatomical measurements

The flies were anesthetized on ice, briefly washed with ethanol and dissected under PBS-T (PBS + 0.1% Triton X-100). Disassembling the fly into manageable elements allowed us to use high magnification, high numerical aperture (NA) objectives that have – in relation to the size of a fly’s body – short working distances but have the benefit of higher axial resolution than the lower magnification/NA objectives. Heads, wings, thoraces with abdomens, fore-, mid- and hind legs were transferred to individual tubes. All body parts except the wings were incubated with 0.25 mg/ml trypsin in PBS-T for 48 hours at 37°C to remove the soft tissues. The cuticle was then bleached in 20% H_2_O_2_ for 24h, and the exoskeleton. The tendons were stained overnight with Congo Red (0.5 mg/mL, Sigma-Aldrich #C676-25G), a bright and comparatively photo-stable chitin-binding dye stains both soft, membranous as well as hard, sclerotized cuticle. It also shows affinity to tendons and fine tendrils, which is very convenient for identification of muscles’ origins and insertion sites, even in the absence of soft tissues. The samples were dehydrated in ethanol and mounted in methyl salicylate (Sigma-Aldrich #M6752), which has refractive index very close to that of glass, facilitated imaging throughout the relatively thick/bulky samples without degradation of the signal. Serial optical sections were obtained on a Zeiss 880 confocal microscope at 2 µm with a Plan-Apochromat 10x/0.45 NA objective, 1 µm intervals with a LD-LCI 25x/0.8 NA objective, or 0.3 µm with a Plan-Apochromat 40x/1.3 NA objective. The 560 nm laser line was used to excite Congo Red.

### Blender model of body geometry

3D meshes were extracted from the confocal stacks using Fiji’s 3D viewer plugin [Schindelin et al., 2012] and imported into Blender [Community, 2018]. A 3D model was constructed from meshes representing the head, thorax+abdomen, wing and fore-, mid- and hind-leg of a single female fly. Appendage meshes were mirrored across the body’s medial plane (Extended Data Fig. 1a). This model was used as the reference for creating a lower polygon count, simplified one, in which the total number of vertices was reduced from 22.6M to 20K (Extended Data Fig. 1b). This simplified model consisted of 67 articulated body segments (Extended Data Fig. 1d): 9 body axis segments (head, thorax, 7 abdominal segments), proboscis (4 segments), antennae, wings, halteres (6 total) and legs (coxa, femur, tibia, 4 tarsal segments and tarsal claws, 6× 8 segments). The exact positions of joints, articulations, and axes of joints’ rotation were determined with high confidence from confocal microscopy data (Fig. 1b, Extended Data Fig. 1c). The model was posed in the rest position and rigged in Blender by creating constraints defining movement of the body segments with respect to each other. Each of the 67 body segments was assigned (parented to) a control element called “bone”, forming a hierarchical kinematic tree system resembling a skeleton called “armature” (Extended Data Fig. 1d).

### MuJoCo model of body physics

The Blender model was then exported to MuJoCo using the dm_control exporter. The components representing head, thorax, abdomen, wings and legs were assigned densities based on average values from weighing 2 groups of 30 and 22 disassembled wild type female flies: head - 0.15 mg, thorax - 0.34 mg, abdomen - 0.38 mg, legs (each) - 0.0162 mg, wings (each) - 0.008 mg. This corresponds to total fly mass of 0.983 mg. The full body length of the model is 0.297 cm, wing span 0.604 cm.

**Extended Data Figure 1:**
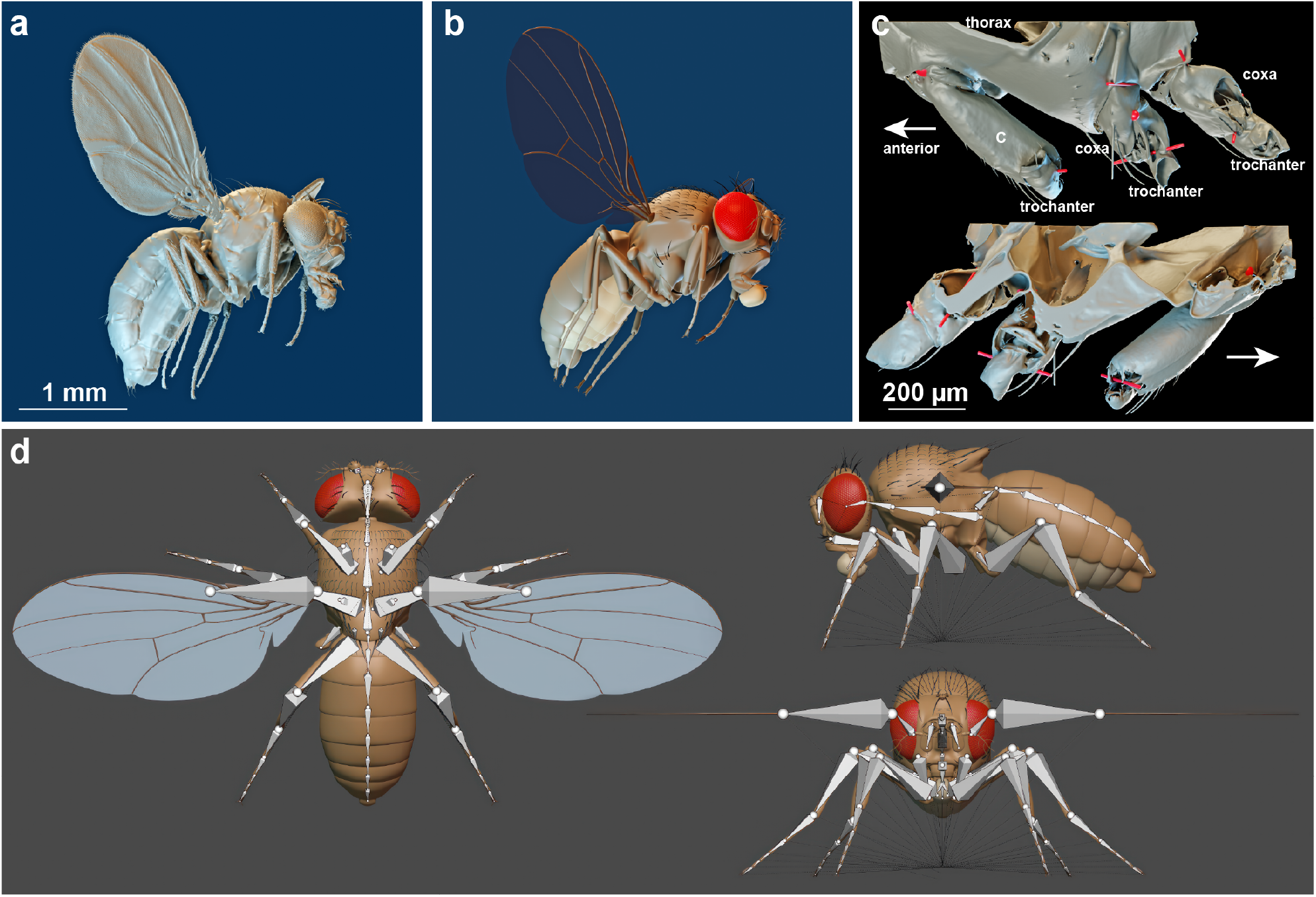
Constructing a 3d model of a female fruit fly from confocal data. (**a**) Hi-polygon (∼22.6 M faces) model of the fly reconstructed in Blender. (**b**) Simplified model (∼20 K faces). (**c**) 3D model of the left-side coxae based on the confocal data. Red bars represent hinge joints, spheres represent ball joints. Arrows indicate anterior. (**d**) Dorsal, sagittal and frontal views of the rigged model in the rest position, the elements of the armature called “bones” shown as elongated octahedrons.

Joint limits were initially determined using Blender’s inverse kinematics tool. We started with fairly tight joint limits and then used reference images of extreme articulated postures (mostly from grooming behaviors) to increase joint limits as required, until all reference poses could be achieved. We then refined the leg joint limits using automated inverse kinematics fitting of the model to 392 frames from manually annotated grooming behavior videos, see below. The sensory system details in the model’s default configuration is shown in Table 2. Degrees of freedom were actuated using torque or position actuators, with certain DoFs (abdomen, tarsi) coupled by tendons (Table 3). For position actuators, control ranges were set to be equal to the corresponding joint ranges. For more detail on building the fly MuJoCo model, see Supplement. For task-specific model modifications, see the Methods sections on Flight and Walking below.

**Table 3:** Actuators of the fly model in its default configuration. Task-specific model modifications (e.g., disabling legs for flight) may include removing some of the actuators. By default, each actuator actuates one DoF. In the “position coupled” category, one actuator actuates several DoFs coupled by a MuJoCo tendon.

| Actuator category | Located at | # of actuators |
| --- | --- | --- |
| torque | wings | 6 |
| position | neck | 3 |
|  | proboscis | 5 |
|  | antennae | 6 |
|  | legs (excluding tarsi) | 42 |
| position coupled<br>(several DoFs per actuator) | tarsi | 6 |
|  | abdomen | 2 |
| adhesion | leg tips | 6 |
|  | labrum | 2 |
| total |  | 78 |

### Analysis of leg degrees of freedom

In order to verify our approximation of the leg degrees of freedom and leg joint ranges we applied the following procedure. We recorded 2-camera videos [Williamson et al., 2018] of multiple free *Drosophila* during grooming behavior. We then uniformly sampled and annotated individual frames of the fly postures during grooming, giving us 3D coordinates of five keypoints for each leg: the four leg joints (body-coxa, coxa-femur, femur-tibia, tibia-tarsus), plus the tarsal tip. We annotated all six legs per frame regardless which legs were actively involved in grooming in the frame. This provided us with data for legs both in grooming positions and in rest (standing) positions. We only observed grooming with T1 and T3 legs and we collected a total of 392 frame annotations. Then we performed inverse kinematics fitting of the model legs to the annotated frames as follows (for details on the inverse kinematics fitting procedure, see also the walking reference data preparation below.) In order to decouple the effect of fly-to-fly size/proportion variability and the actual degrees-of-freedom mismatch, in each frame we rescaled the model’s leg segments to match data. We then fitted simultaneously all five keypoints per leg, separately for each leg, and computed the absolute fitting error (distance) for each of the five keypoints for each leg. Extended Data Fig. 2 shows the distributions of the inverse kinematics fitting errors for each keypoint and each leg. Extended Data Fig. 2a shows the errors for leg fits in rest position, Extended Data Fig. 2b shows errors in grooming positions. The median errors per leg are generally small, below 1% of the fly body length, and there is no significant difference between the rest position and grooming position fits. There appears to be a slight systematic increase in the tibia-tarsus keypoint error, more noticeable in the grooming fits in Extended Data Fig. 2b, which is not surprising as grooming leg positions tend to be more intricate than the rest position. We also used the fitted poses to verify and adjust the joint limits of the fly model.

### Distributed Reinforcement Learning

For each locomotion task, we trained a policy network using a distributed RL setup [Nair et al., 2015, Horgan et al., 2018] powered by Ray, an open-source general-purpose distributed computing package [Moritz et al., 2017]. The distributed training configuration is shown in Extended Data Fig. 3. Multiple CPU-based actors run in parallel in separate MuJoCo environment instances, generate experiences, and log them into a replay buffer. A single GPU-based learner samples training batches from the replay buffer and updates the policy and critic network weights. The critic network is a part of the training process only and is not used by the fly model directly. Each actor explores the environment and generate experiences using its own copy of the policy network whose weights are periodically synchronized with the current learner policy. For learner policy updates, we used the off-policy actor-critic DMPO agent, a distributional extension [Bellemare et al., 2017] of the MPO agent [Abdolmaleki et al., 2018b, Abdolmaleki et al., 2018a]. We used `dm_control` [Tunyasuvunakool et al., 2020] to set up the RL environments and for MuJoCo python bindings. We used the DMPO agent implemented in `acme` [Hoffman et al., 2020] and the replay buffer implemented in `reverb` [Cassirer et al., 2021], and the `Adam` optimizer [Kingma and Ba, 2017]. The hyperparameters of the distributed setup and of the DMPO agent are shown in Table 4 and in Table 5.

**Table 4:** Distributed training hyperparameters. See implementation for more detail.

| Hyperparameter | Value (walking) | Value (flight) |
| --- | --- | --- |
| num_actors | 32 | 32 |
| batch_size | 256 | 256 |
| prefetch_size | 4 | 4 |
| min_replay_size | $10^4$ | $10^4$ |
| max_replay_size | $4 \times 10^6$ | $4 \times 10^6$ |
| samples_per_insert | 15 | 15 |
| n_step | 10 | 5 |
| num_samples | 20 | 20 |
| policy_optimizer_lr | $10^{-4}$ | $10^{-4}$ |
| critic_optimizer_lr | $10^{-4}$ | $10^{-4}$ |
| dual_optimizer_lr | $10^{-3}$ | $10^{-3}$ |
| target_critic_update_period | 200 | 107 |
| target_policy_update_period | 200 | 101 |

**Table 5:** DMPO agent hyperparameters. See implementation for more detail.

| Hyperparameter | Value (walking) | Value (flight) |
| --- | --- | --- |
| epsilon | 0.05 | 0.1 |
| epsilon_mean | $5 \times 10^{-5}$ | $2.5 \times 10^{-3}$ |
| epsilon_stddev | $10^{-7}$ | $10^{-7}$ |
| action_penalization | False | True |
| epsilon_penalty | N/A | 0.1 |
| init_log_temperature | 10 | 10 |
| init_log_alpha_mean | 10 | 10 |
| init_log_alpha_stddev | $10^3$ | $10^3$ |
| per_dim_constraining | True | True |

To guarantee stability, we ran MuJoCo physics simulations at ×4–10 smaller time steps than we sampled action commands from the policies [Tunyasuvunakool et al., 2020], see the physics and control time-step values in Tables 10, 14, 18. The policies were stochastic during training (outputting distribution over actions) to facilitate exploration. The distributions were Gaussian, independent for each action dimension and parameterized by mean and standard deviation. The policy network architectures for each task are given in Tables 11, 15, 19. At test time, the policies reverted to being deterministic by using the means of the predicted action distributions. The actions output by the policy networks were in the canonical range [− 1, 1]. The actions were then rescaled to match their corresponding proper ranges in the fly model, e.g. joint limits for position actuators or force limits for force actuators. All observables (policy inputs) were strictly egocentric, e.g. calculated in the local reference frame of the fly model. We only used feedforward policy and critic networks and did not extensively sweep the network architectures.

**Extended Data Figure 2:**
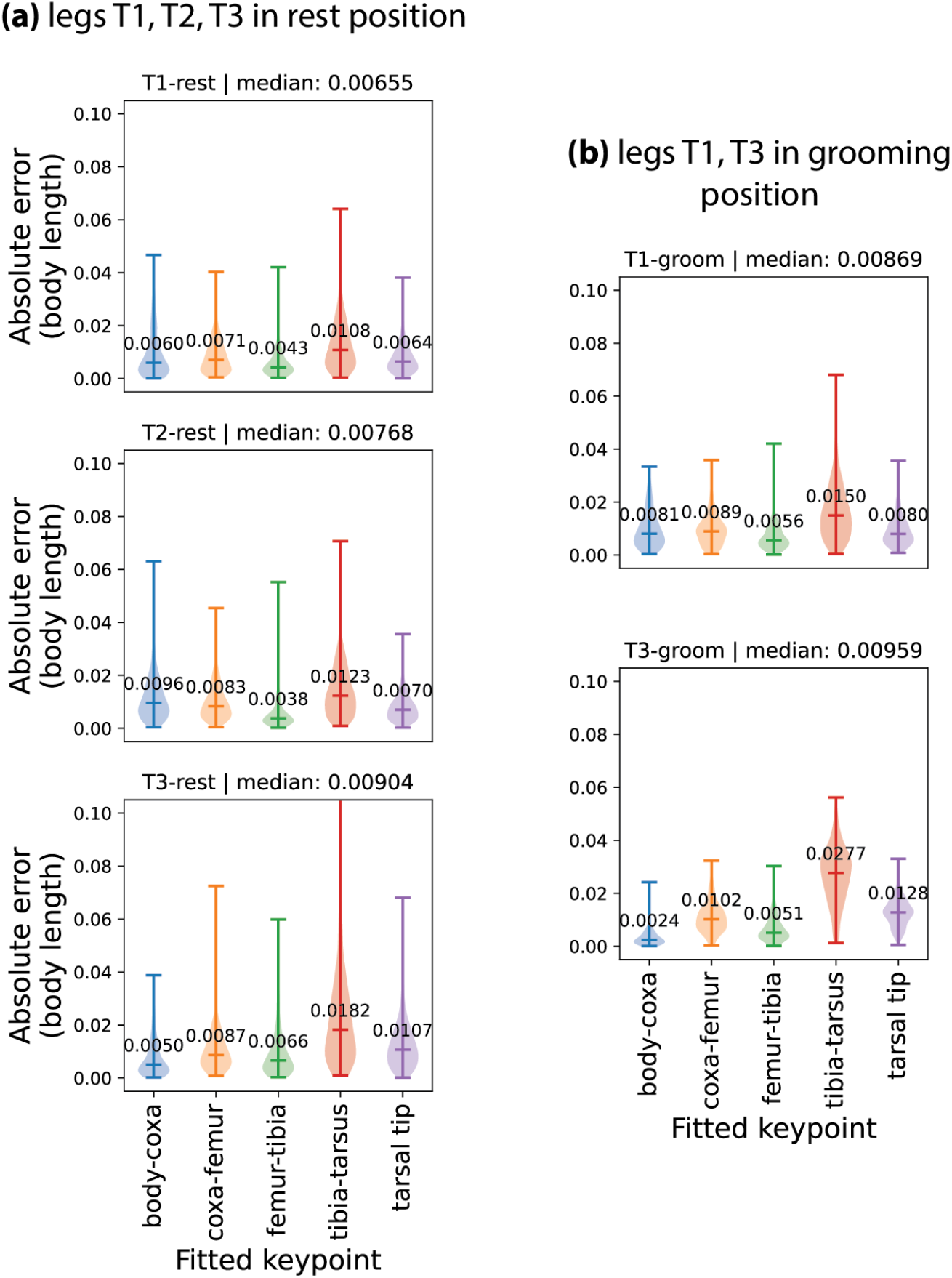
Leg degrees of freedom analysis. Inverse kinematics fits of the fly model legs to real *Drosophila* grooming poses (392 poses in total.) To separate the effect of degrees-of-freedom from fly-to-fly size variability, model legs were rescaled to match these in each individual reference pose frame. Individual model legs were fitted separately by simultaneously matching five leg keypoints located at four leg joints and leg tip. Absolute fitting errors for each leg keypoint are shown for (**a**) all leg pairs in rest position, (**b**) leg pairs T1 and T3 in grooming positions. Horizontal bars and corresponding values are median errors for each keypoint. Median errors across all keypoints in each leg pair are also indicated.

**Extended Data Figure 3:**
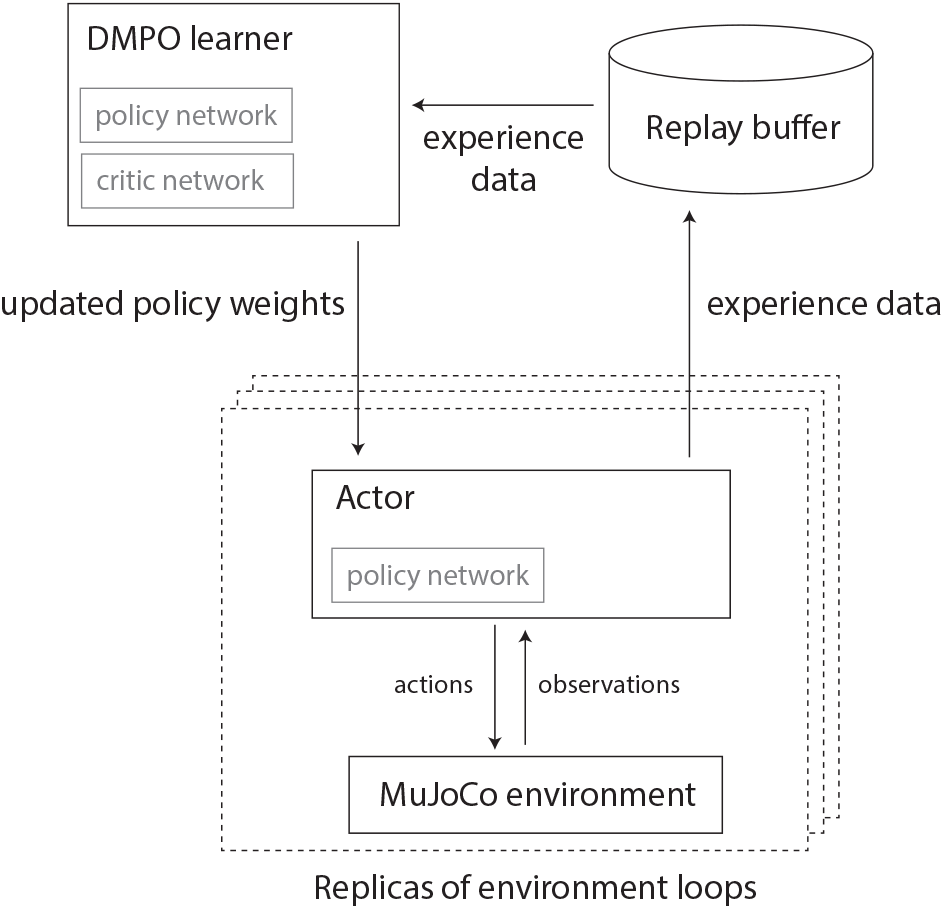
Distributed training architecture. Multiple replicas of actors in MuJoCo environments collect experiences and feed them to a single replay buffer. The DMPO learner samples experiences from the replay buffer, updates the policy and critic network weights, and sends the updated weights to the actors’ copies of the policy.

We trained the model in episodes of finite number of time steps. An episode ends either when (i) the episode time limit is successfully reached, or (ii) when an early termination condition, signifying failure, is met. Termination condition details are explained in the corresponding RL task sections below. In the first case, the agent estimates the remaining infinite-horizon future rewards (beyond episode’s final step) by bootstrapping from the value of the state at the end of the episode. In the second case, the reward sequence is truncated by setting the future reward to zero. The loss of the future infinite-horizon rewards is an unfavorable outcome and the agent will try to learn to avoid events triggering early episode termination.

### Phenomenological model of the fluid forces

This section describes the fluid force model we introduced to MuJoCo to facilitate the simulation of fly’s flight. The fluid model computes forces exerted on moving rigid bodies whose shape can be approximated by ellipsoids. The model provides fine-grained control of the different types of fluid forces via five dimensionless coefficients, the `fluidcoef` attribute in MuJoCo, see the MuJoCo documentation for more detail ^1^. We used the fluid model to compute forces on fly’s flapping wings whose shape closely matches slender ellipsoids (Fig. 1g). Elements of the fluid force model are a generalization of [Andersen et al., 2005] and [Berman and Wang, 2007] to three dimensions. The force **f** and torque **g** that the surrounding fluid exerts onto the translating and rotating body are approximated as a sum of effects: a lifting force due to the translational circulation *f*_*K*_, an additional lifting force due to the rotational circulation of the body *f*_*M*_, a force and torque due to the viscous drag **f**_*D*_ and **g**_*D*_, and finally the effect of the added mass **f**_*A*_ and **g**_*A*_:

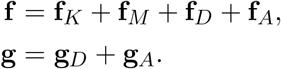

The MuJoCo model is implemented generally, and they are referred to respectively as Kutta lift, Magnus force, viscous drag and resistance and added mass. We will user this naming convention in the section below. The forces imposed by the fluid onto each ellipsoid are computed independently by approximating the effect of an incompressible quiescent fluid of density *ρ* and (dynamic) viscosity *ν*. The problem is described in a reference frame aligned with the principal axes of the ellipsoid and moving with it. The ellipsoid has semi-axes **r** = {*r*_*x*_, *r*_*y*_, *r*_*z*_}, volume *V* = (4*π/*3)*r*_*x*_*r*_*y*_*r*_*z*_, velocity **v** = {*v*_*x*_, *v*_*y*_, *v*_*z*_}, and angular velocity ***ω*** = {*ω*_*x*_, *ω*_*y*_, *ω*_*z*_}. We will also use *r*_max_ = max {*r*_*x*_, *r*_*y*_, *r*_*z*_, *r*_min_} = min {*r*_*x*_, *r*_*y*_, *r*_*z*_}, and *r*_mid_ = *r*_*x*_ + *r*_*y*_ + *r*_*z*_− *r*_max_ −*r*_min_. The area projected by the ellipsoid onto the plane normal to **v** is

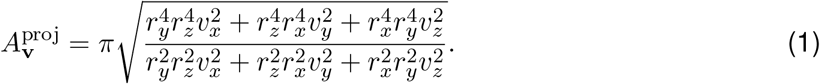

The circulation Γ is the line integral of the fluid velocity field **v**_*f*_ around a closed curve Γ = ∮ **v**_*f*_ · d**l**. The individual force and torque components are described below.

#### Kutta lift

The Kutta condition describes an effect that is valid also for potential (i.e. inviscid) flows. For a body moving in a potential flow there are two stagnation points (a location in the flow field where the velocity is zero): in the front, where the stream-lines separate to either sides of the body, and in the rear, where they reconnect. The Kutta condition is the observation that a moving body with a sharp rear edge will generate in the surrounding flow a circulation of sufficient strength to hold the rear stagnation point at the trailing edge. In two-dimensional potential flow, the circulation due to the Kutta condition for a slender body can be estimated as Γ_K_ = *C*_*K*_ *r*_*x*_ ∥**v**∥ sin 2*α*. Here *C*_*K*_ is a Kutta lift coefficient and *α* is the angle between the velocity vector and its projection onto the surface. The lift force per unit length can be computed with the Kutta–Joukowski theorem as **f**_*K*_*/ℓ* = *ρ*Γ_K_ × **v**.

**Extended Data Figure 4:**
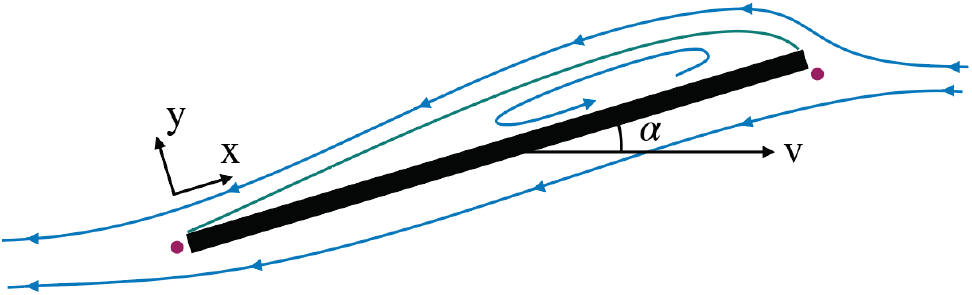
Schematic of the Kutta condition. Blue lines are streamlines and the two magenta points are the stagnation points. The dividing streamline, which connects the two stagnation points, is marked in green. The dividing streamline and the body inscribe an area where the flow is said to be “separated” and recirculates within. This circulation produces an upward force acting on the plate.

In order to extend the lift force equation to three-dimensional motions, we consider the normal 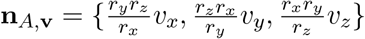 to the cross-section of the body which generates the body’s projection 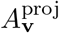 onto a plane normal to the velocity. We use this direction to decompose **v** = **v**_∥_ + **v**_⊥_ with 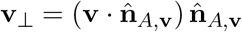 (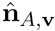 is the unit vector). We write the lift force as:

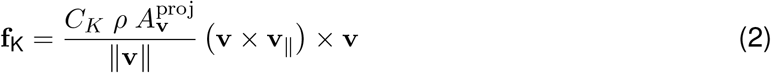

Note that the direction of 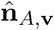 differs from **v** only on the planes where the semi-axes of the body are unequal. So for example, for spherical bodies 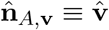 and by construction **f**_K_ = 0.

#### Magnus force

A spinning body induces rotation in the surrounding fluid which deflects the trajectory of the fluid flow past the body, and the body receives an equal an opposite reaction. Following [Andersen et al., 2005], we estimate the force due to the rotation as

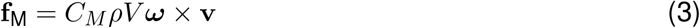

where *V* is the volume of the body and *C*_*M*_ is a coefficient for the force, which we typically set to 1.

#### Viscous drag

The drag force acts to oppose the motion of the body relative to the surrounding flow. For high Reynolds numbers, the viscous drag can be approximated with the drag equations, first proposed by Newton, *f*_D_ = − *C*_*D*_*ρs*_*D*_*v*^2^ and *g*_D_ = − *C*_*D*_*ρI*_*D*_*ω*^2^. Here *C*_*D*_ is a drag coefficient, *s*_*D*_ is a reference surface area (e.g., a measure of the area projected on the plane normal to the flow), and *I*_*D*_ is a reference moment of inertia. These quantities depend on the properties of the fluid, the shape of the body and its velocity [Duan et al., 2015]. We derive a correlation for the viscous drag based on two surfaces: the projection surface 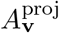 of the ellipsoid onto a plane normal to **v** and the maximum projected surface *A*_max_= *πr* _*max*_ *r*_*mid*,_, such that 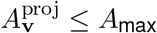:

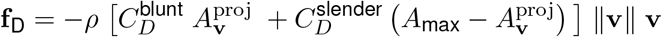

We propose an analogous model for the angular drag. For each Cartesian axis we consider the moment of inertia of the maximum swept ellipsoid obtained by the rotation of the body around the axis. The resulting components of the moment of inertia are:

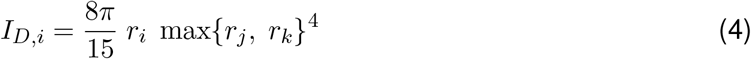

where, as before, the indices *i, j, k* are cyclic permutations of the axes (*x, y, z*), (*y, z, x*), (*z, x, y*). The angular drag torque is computed as:

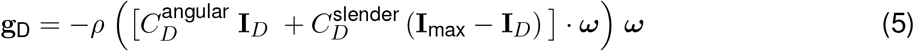

Here **I**_max_ is a vector with each entry equal to the maximal component of **I**_*D*_.

For low Reynolds numbers, the viscous drag is well approximated by Stokes’ law [Stokes, 1850] for an equivalent sphere:

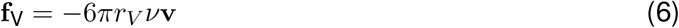

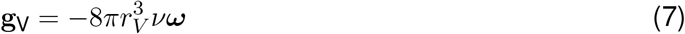

where *r*_*V*_ = (*r*_*x*_ + *r*_*y*_ + *r*_*z*_)*/*3 is the radius of the equivalent sphere. We add this term to the previously defined viscous drag force and torques **f**_*D*_ and **g**_*D*_ to approximate the drag well for both high and low Reynolds numbers.

#### Added mass

Added mass measures the inertia of the fluid that is put into motion by the body’s motion. In the case of a body with three planes of symmetry, the forces **f**_*A*_ and torques **g**_*A*_ can be written as [Lamb, 1932]:

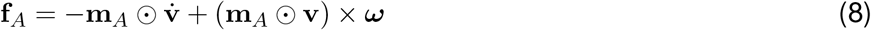

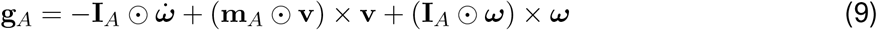

Here, ⊙ denotes an element-wise product, 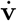 is the linear acceleration and 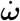 is the angular acceleration. **m**_*A*_ ⊙ **v** and **I**_*A*_ ⊙***ω*** are the virtual linear and angular momentum respectively. The added-mass vector **m**_*A*_ = {*m*_*A,x*_, *m*_*A,y*_, *m*_*A,z*_} and added-moment of inertia vector **I**_*A*_ = {*I*_*A,x*_, *I*_*A,y*_, *I*_*A,z*_} measure the inertia of the fluid displaced by the motion of the body in the corresponding direction and can be derived from potential flow theory for certain simple geometries.

For an ellipsoid, the virtual mass *m*_*A,i*_ for a motion along axis *i* and the virtual moment of inertia *I*_*A,i*_ for a rotation along axis *i* are:

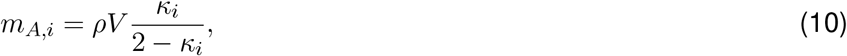

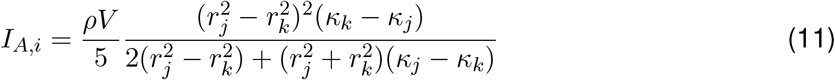

where the dimensionless virtual inertia coefficients are [Tuckerman, 1925]:

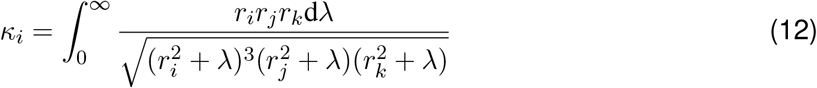

which we compute by 15-point Gauss–Kronrod quadrature. The indices *i, j, k* are cyclic permutations of the axes (*x, y, z*), (*y, z, x*), (*z, x, y*).

### Modeling flight behavior

#### Wing actuator gain, wing joint damping, MuJoCo fluid model parameters

We used the following procedure to fit the flight physics parameters. We started with a wing motion trajectory recorded previously from a hovering *Drosophila Melanogaster* [Dickson et al., 2008] ^2^. We placed the model in a hovering position and actuated the wings to reproduce the real wing trajectory by using the real wing angles as target angles for the wing actuators. We then iteratively adjusted (increased) the wing actuator gain to a point where the mean absolute error between the reference wing angles and the trajectory traversed by the model’s wings was below 5% of the wing angle amplitude. At each iteration, we also fitted the wing joint damping coefficient to avoid underdamping and ensuing wing oscillations. We used the same gain value for all three wing actuators (yaw, roll, pitch). Our final values for the gain and damping pair were `gainprm = [18, 18, 18], damping = 0.007769.`

Having found suitable wing actuator gain and wing joint damping, we adjusted the MuJoCo fluid model coefficients (Table 6) which scale the drag and lift forces produced by the flapping wings. These (dimensionless) fluid model coefficients are defined in the fluid model Methods section and stored in the `fluidcoef` MuJoCo attribute. We placed the model in a flight position and again drove the wings with the real reference angles as target angles for the wing actuators. We then iteratively found a set of fluid parameters such that the net lift approximately balanced the fly model weight during several wingbeat cycles, `fluidcoef = [1.0, 0.5, 1.5, 1.7, 1.0].` The flight physics parameters are summarized in Table 7.

**Table 6:** Fluid force model coefficients. See Section for definition. The coefficients are stored in the `fluidcoef` MuJoCo attribute, in this order.

| Index | Coefficient | Description | Value |
| --- | --- | --- | --- |
| 0 | $C_D^{\text{blunt}}$ | Blunt drag | 1.0 |
| 1 | $C_D^{\text{slender}}$ | Slender drag | 0.5 |
| 2 | $C_D^{\text{angular}}$ | Angular drag | 1.5 |
| 3 | $C_K$ | Kutta lift | 1.7 |
| 4 | $C_M$ | Magnus lift | 1.0 |

**Table 7:** Flight physics parameters.

| Parameter | Value | Units |
| --- | --- | --- |
| `fluidcoef` | [1.0, 0.5, 1.5, 1.7, 1.0] | unitless |
| `gainprm` | [18, 18, 18] | dyn cm |
| `damping` | 0.007769 | dyn s |
| `body_pitch_angle` | 47.5 | deg |
| `stroke_plane_angle` | 0 | deg |
| `base_freq` | 218 | Hz |

We also performed a sensitivity analysis on the the viscous drag and Kutta lift, the two dominant forces in the fluid dynamics. We re-trained imitation learning of free flight with modified choices for the coefficients associated with viscous drag and Kutta lift, with all other coefficients held fixed. We then evaluated the degree to which imitation learning was able to correctly reproduce ground truth center of mass flight trajectories with realistic wing kinematics, as in the original experiments reported in the paper. We also quantified the fraction of trajectories where the fly crashed to the ground as a second performance measure. Figure 5 shows that flight performance is robust to even 20% variation in these parameters.

#### Wingbeat Pattern Generator

All our flight tasks utilize a wingbeat pattern generator (WPG) that produces, in an open-loop fashion, a fixed mirror-symmetric cyclic baseline wing trajectory. The WPG generates the baseline pattern by design with no learning involved. The baseline trajectory closely follows a previously recorded wing pattern of a hovering *Drosophila Melanogaster* [Dickson et al., 2008]. The baseline pattern is available at Supp. Data. At each simulation time-step, the WPG retrieves and outputs the six wing angles (three per wing, Fig. 2) of the baseline wing pattern for the current wing beat cycle step. These baseline wing angles get converted to torque action commands for the wing actuators. While already producing a realistically looking wing motion, the fixed baseline wing trajectory alone is not sufficient to support a stable hovering due to the lack of feedback loop, approximations in the MuJoCo fluid model, and the sim-to-real gap. It is the role of the policy network to provide these missing components. To achieve this, the WPG torque action is combined (additively) with the policy output to produce the final action vector to be sent to the wing actuators. In this way, the policy modulates the fixed baseline pattern and produces flight required by the task at hand, e.g. stabilize flight, hover, speed up, turn, etc.

**Extended Data Figure 5:**
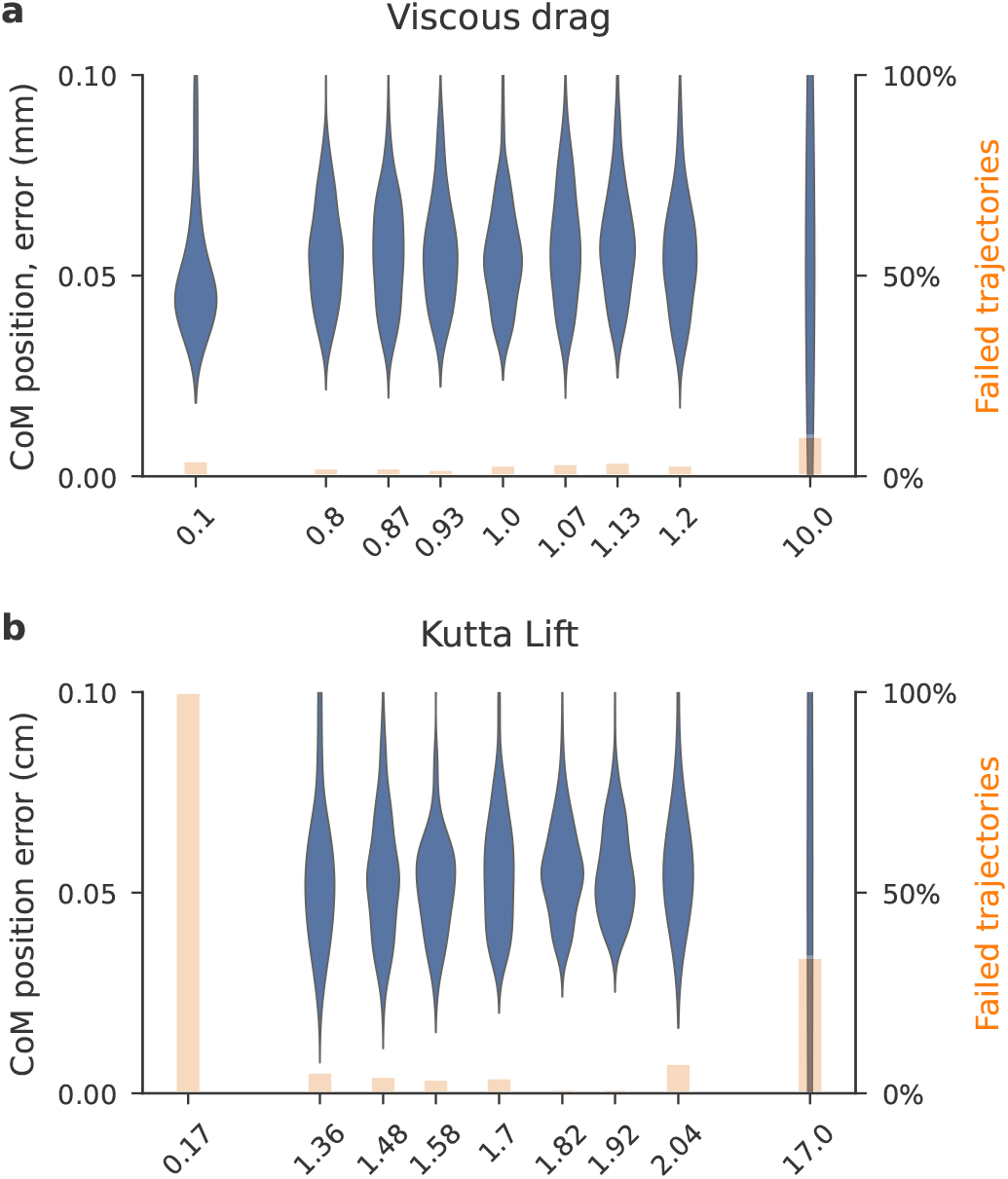
Sensitivity analysis of the dimensionless fluid model scaling coefficients for viscous drag (a) and Kutta lift (b) components. Accuracy of imitation learning is shown in blue violins (body center of mass error, 5 training runs, same trajectories as in 2) and trajectory failure rate in orange bars (percentage of trajectories which crashed) for simulations varying each coefficient by up to 10x.

The wing motion produced by the combination of the WPG and the trained policy stays close to the initial WPG baseline pattern. This is achieved by penalizing the magnitude of the policy actions during training with the DMPO agent [Abdolmaleki et al., 2018b, Abdolmaleki et al., 2018a]. In this way, the agent is encouraged to discover a physically viable wing motion pattern by using only minimal policy actions without deviating significantly from the baseline pattern. In addition, the WPG can vary the frequency of the output baseline pattern within a pre-defined range. A single scalar out of the policy action vector is used by the WPG to control the baseline wingbeat frequency. In our setting, the wingbeat frequency was allowed to vary within 10% range centered at 218 Hz, *Drosophila melanogaster* average frequency [Fry et al., 2005]. The WPG is implemented as a lookup table containing the single fixed baseline wing pattern resampled at wingbeat frequencies within the 10% frequency range. When a frequency change is requested by the policy, the WPG will smoothly connect the patterns at old and new frequencies.

#### Flight imitation task configuration

As the flight reference data, we used previously recorded trajectories of freely flying *Drosophila hydei*. The trajectories contain a fly’s Cartesian center-of-mass position and body orientation represented as a quaternion. The trajectories were recorded at 7500 fps. We started with 44 trajectories of spontaneous turns (saccades) [Muijres et al., 2015] and 92 trajectories of evasion maneuvers [Muijres et al., 2014] in response to visual looming stimuli. Each reference trajectory started with the fly first flying normally and then performing a maneuver. During and after the maneuver, the fly could fly straight, sideways, and backwards. The flies could also ascend and descend. We linearly interpolated the raw trajectories to the flight simulation control step of 0.2 ms. Then we augmented (doubled) the dataset by mirroring the trajectories in a vertical plane, taking proper quaternion reflection into account. This resulted in a dataset of 272 flight trajectories, equivalent to ∼ 53 seconds of real time flight. The dataset is available at Supp. Data. We used 80% of the trajectories for training and the rest for testing. Due to the small size of the dataset, to maintain balance between left and right turns in the training data, we split the dataset such that if a trajectory is in the training set, so is its mirrored counterpart. We simulated flight at 0.05 ms physics time-steps and 0.2 ms control time-steps, Table 10.

The reinforcement learning task is set as follows. In each episode, the fly model is required to track a reference trajectory selected from the flight dataset at random. The episode begins from a random step within the selected reference trajectory, excluding the last 50 steps. The model’s initial position, orientation, linear and rotational velocities are set equal to the reference. The initial phase of the wing cycle is randomized. The episode ends either when the end of trajectory is successfully reached, or terminates early if the model hits the ground or is displaced from the reference CoM position by more than 2 cm.

The reward is calculated based on how closely the fly model tracks the reference trajectory at each timestep. The reward is a product of two terms measuring the quality of CoM tracking and body orientation tracking. At every simulation timestep, the reward *R* ∈ [0, 1] is computed as:

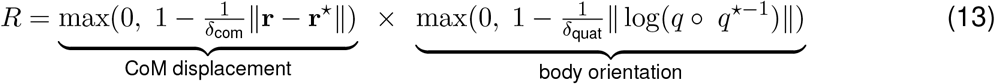

The first term measures the current displacement between the model and reference CoM positions, **r** and **r**^⋆^ respectively. The second term is the body orientation mismatch, calculated as the norm of the quaternion “minus operator” between the model and reference quaternions, *q* and *q*^⋆^, respectively [Sola, 2017]. *δ*_com_ and *δ*_quat_ are the reward tightness hyperparameters. For instance, the reward is zero when the CoM displacement in the first term ∥**r** −**r**^⋆^ ∥ *>δ*_com_, while it is one when ∥**r**− **r**^⋆^ ∥= 0, and similarly for the second term. See Table 10 for the hyperparameter values used.

In the flight imitation task, we placed the fly legs in a retracted flight position and disabled the leg DoFs and actuators. The retracted leg configuration is stored in the `springref` leg parameters, which were fitted by matching the model’s legs to images of flying *Drosophila*. We also removed the antennae and proboscis actuators and excluded their joints from the observations. This reduced the number of observable joint angles and joint velocities to 25. It also reduced the total action dimension to 12. We didn’t use vision in this task. The observables (policy inputs) are listed in Table 8 and the actions (policy outputs) are shown in Table 9. In addition to the standard set of egocentric vestibular and proprioception observables, the policy receives task-specific inputs: the Cartesian CoM displacement and the orientation (quaternion) displacement of the reference trajectory with respect to the model at the current timestep plus 5 timesteps into the future. The displacements are calculated with respect to the current fly model position and orientation and are expressed in the egocentric reference frame of the model. The policy network architecture is shown in Table 11. The trained flight imitation policy can be used as a low-level flight controller. In this scenario, the two task-specific inputs (CoM and quaternion displacements) serve as high-level steering control commands.

**Table 8:** Observations in the flight imitation task. All observables are calculated in fly’s egocentric reference frame.

| Category | Observables (policy inputs) | Units | Array shape |
| --- | --- | --- | --- |
| vestibular | velocimeter | cm/s | 3 |
|  | accelerometer | cm/s<sup>2</sup> | 3 |
|  | gyro | rad/s | 3 |
|  | gravity direction | cm | 3 |
| proprioception | joint angles | rad | 25 |
|  | joint velocities | rad/s | 25 |
| task inputs<br>(steering commands) | reference CoM displacement | cm | (6, 3) |
|  | reference orientation displacement | unitless quaternion | (6, 4) |
| total obs. dimension |  |  | 104 |

**Table 9:** Actions in the flight imitation task.

| Actions (policy outputs) | Array shape |
| --- | --- |
| head | 3 |
| wings | 6 |
| abdomen | 2 |
| WPG frequency control | 1 |
| total action dimension | 12 |

**Table 10:**
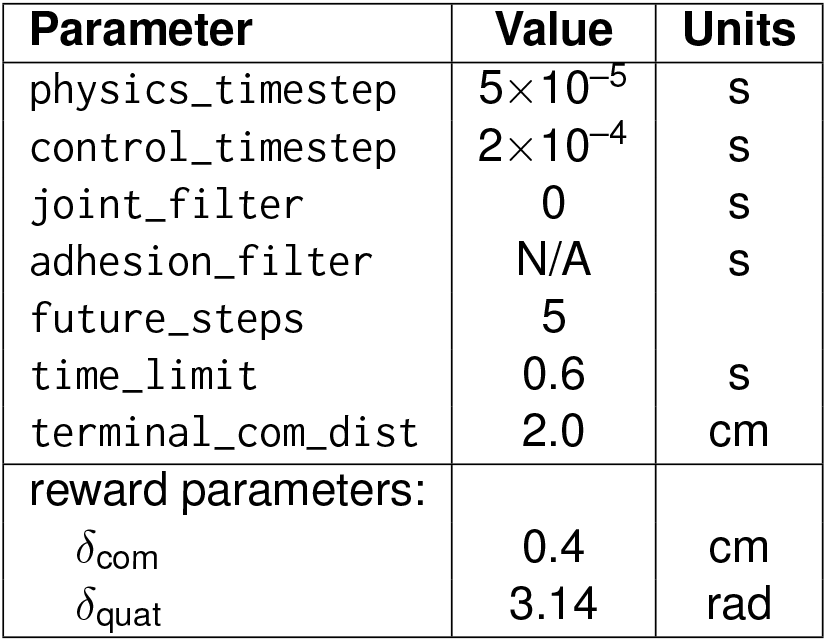
Flight imitation task hyperparameters. See implementation for more details.

| Parameter | Value | Units |
| --- | --- | --- |
| physics_timestep | $5 \times 10^{-5}$ | s |
| control_timestep | $2 \times 10^{-4}$ | s |
| joint_filter | 0 | s |
| adhesion_filter | N/A | s |
| future_steps | 5 |  |
| time_limit | 0.6 | s |
| terminal_com_dist | 2.0 | cm |
| reward parameters: |  |  |
| $\delta_{\text{com}}$ | 0.4 | cm |
| $\delta_{\text{quat}}$ | 3.14 | rad |

**Table 11:** Flight policy network architecture. Layer sizes indicated in parentheses.

|  |  |
| --- | --- |
| *input: observations* |  |
| Linear (256) |  |
| LayerNorm |  |
| tanh |  |
| Linear (256) |  |
| ELU |  |
| Linear (256) |  |
| ELU |  |
| Linear (12) | Linear (12) |
|  | softplus |
| *output: action mean* | *output: action std* |

At training time, we engage action penalization, a DMPO agent feature. It encourages the agent to learn to utilize actions of small magnitude thus keeping the wing motion close to the baseline wing pattern produced by the WPG. The task and agent hyperparameters, including DMPO action penalization, are summarized in Table 5 and Table 10.

#### Flight controller reuse, vision-guided flight task configuration

We reused the flight controller trained in the flight imitation task described above in the context of the two vision-guided reinforcement learning tasks set as follows. The fly model is required to fly generally in the positive *x*-direction, to maintain a given target speed and height above terrain, and to avoid collisions with the terrain. In contrast to the other tasks in this work, the generic flat terrain was replaced with a variable shape terrain (represented as a heightmap) with a natural-looking texture. As the policy has no direct access to the flight height and terrain shape, avoiding collisions and assessing the current height requires utilization of the visual input from the two eye cameras. In each episode, the terrain shape is procedurally regenerated, and a new target speed and target height are randomly selected (Table 14). The episodes end normally when the time limit is reached, or terminate early when the fly collides with terrain, which results in the loss of all future infinite-horizon rewards.

The task is set in a square 40 ×40 cm arena. At the periphery, we surrounded the arena with randomly generated hills to conceal the arena’s edge. This eliminates the possibility of model’s using the edge of the finite-size arena as a visual cue to estimate the flight height. Also, we removed overhead light sources to eliminate the fly’s shadows as the shadow size could be exploited to infer the flight height. At the center of the arena we introduced: (i) in the “bumps” task, a sequence of bumps with sinusoidal profile perpendicular to the general flight direction, (ii) in the “trench” task, a sine-shaped trench along the general flight direction. In both tasks, the amplitude, phase, period, height of the bumps and trench were randomly selected in each episode (see Table 14 for the terrain randomization details). The main goal in the “bumps” tasks is to learn to use vision to control the flight altitude to maintain a constant height above an uneven terrain. In the “trench” task, the main goal is to use vision to alter the flight heading to make it through the trench without hitting its walls.

In the “bumps” tasks, the (multiplicative) reward consists of several factors as follows:

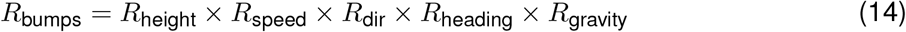

where the individual reward terms are:

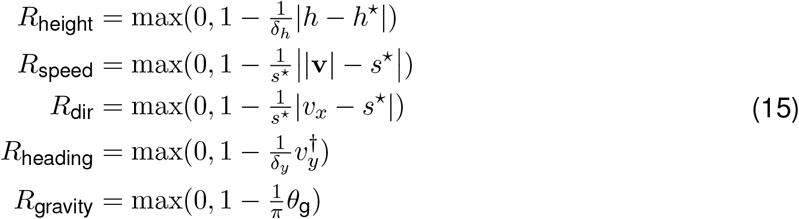

Here, *R*_height_ is the reward factor measuring how close the current flight height *h* is to the target height *h*^⋆^. As in the other tasks, *δ*_*h*_ is the reward tightness hyperparameter (see Table 14). Similarly, *R*_speed_ compares the current flight speed |**v**| to the target speed *s*^⋆^ (**v** is the fly velocity vector). *R*_dir_ prescribes a preferred general flight direction by rewarding fly’s propagation in the positive *x*-direction. *v*_*x*_ is the *x*-component of fly velocity computed in the arena reference frame. *R*_heading_ requires the fly to keep the body heading parallel to the current velocity direction (e.g., avoid flying sideways). This is achieved by minimizing the lateral velocity component 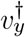 computed in fly’s egocentric reference frame. *R*_gravity_ sets a preferred body orientation with respect to the gravity direction (the vertical *z*-axis) by minimizing *θ*_g_, the angle between the current gravity direction vector **g** as measured by fly’s gravity sensor and the preferred egocentric gravity direction **g**^⋆^.

This angle is computed as *θ*_g_ = cos^−1^(**g** ·**g**^⋆^). The preferred gravity direction is expressed in fly’s egocentric reference frame as **g**^⋆^ = (sin(*α*_pitch_), 0, cos(*α*_pitch_)), where *α*_pitch_ = 47.5^°^ is the default body pitch angle during stable flight in *Drosophila* [Muijres et al., 2014], see Table 7.

In the “trench” task, the reward is as in the “bumps” task with an additional term:

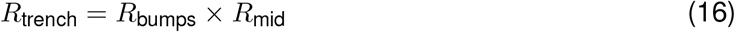

The additional term *R*_mid_ requires the fly to stay close to the midline of the trench and it is written as:

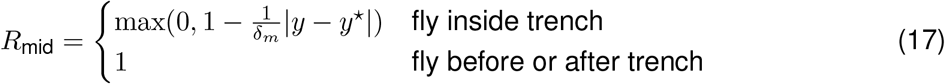

This term compares the current fly’s *y*-coordinate (lateral, perpendicular to the general trench direction *x*) and the current midpoint of the trench, *y*^⋆^. In the “trench” task, the target height is always smaller than the trench wall height. Also, the trench width and turn amplitude are sampled such that the “trivial” solution of making it through the trench by simply flying straight wouldn’t be possible.

In both vision-guided flight tasks, we reused the policy network trained in the flight imitation task (described above) as a low-level flight controller. We froze the weights of this pre-trained low-level controller network and trained a high-level controller to navigate the flight. In this setup, the low-level controller abstracts away and controls all the fine details of the wing motion (Table 13), while the high-level controller only issues low-dimensional steering commands to the low-level controller.

As in the flight imitation task, the fly legs were retracted and their DoFs and actuators disabled. The observations, on the other hand, were altered in two ways. First, we added visual input from the eye cameras as two 32×32 RGB frames. Second, we provided a task-input consisting of the target speed and height, which were randomly sampled in the beginning of each episode. The total observation dimension was 6208, see Table 12 for more detail. The RGB visual input was first converted to grayscale, then processed by a convolutional visual module, and then fed into the high-level controller’s policy network as a vector of dimension 8, along with the rest of the observables. The steering command output by the high-level controller was concatenated with the proprioception and vestibular (but not visual and task-input) components of the original observation and passed on as input to the low-level controller network. We used a generic convolutional network mostly similar to a single block of ResNet[He et al., 2015]. The architecture of the high-level controller network is shown in Table 15. We used the same DMPO agent configuration, including action penalization, as in the flight imitation task, see Table 5. We trained the vision flight controller, including the convolutional visual module, end-to-end with reinforcement learning. The vision task hyperparameters are summarized in Table 14.

**Table 12:** Observations in the vision-guided flight task. All observables are calculated in fly’s egocentric reference frame.

| Category | Observable (policy input) | Units | Array shape |
| --- | --- | --- | --- |
| vision | right eye | RGB (byte) | (32, 32, 3) |
|  | left eye |  | (32, 32, 3) |
| vestibular | velocimeter | cm/s | 3 |
|  | accelerometer | cm/s<sup>2</sup> | 3 |
|  | gyro (rotational velocity) | rad/s | 3 |
|  | gravity direction | cm | 3 |
| proprioception | joint angles | rad | 25 |
|  | joint velocities | rad/s | 25 |
| task input | target flight speed | cm/s | 1 |
|  | target flight height | cm | 1 |
| total observation dimension |  |  | 6208 |

**Table 13:** Actions in the vision-guided flight task. The pre-trained low-level controller outputs the full 12-dimensional actions, same as in the flight imitation task.

| Action (policy output) | Array shape |
| --- | --- |
| head | 3 |
| wings | 6 |
| abdomen | 2 |
| WPG frequency control | 1 |
| total action dimension | 12 |

**Table 14:** Vision-guided flight task parameters and ranges of sine bump and trench terrain randomization.

| Parameter | Value | Units |
| --- | --- | --- |
| eye_camera_fovy | 150 | deg |
| eye_camera_size | 32 | pixels |
| vis_output_dim | 8 |  |
| time_limit | 0.4 | s |
| reward parameters: |  |  |
| $\delta_h$ | 0.15 | cm |
| $s^*$ | $1.1 \times \text{target\_speed}$ | cm/s |
| $\delta_y$ | 10 | cm/s |
| $\delta_m$ | 0.15 | cm |
| flight speed range | [20, 40] | cm/s |
| flight height range | [0.5, 0.8] | cm |
| bump terrain parameters: |  |  |
| period range | [10, 15] | cm |
| height range | [0.5, 1] | cm |
| phase range | [0, $2\pi$] | rad |
| trench terrain parameters: |  |  |
| period range | [5, 8] | cm |
| amplitude range | [0.35, 0.6] | cm |
| phase range | [0, $2\pi$] | rad |
| width range | $2 \times \text{amplitude} + [0.3, 0.6]$ | cm |
| height | 1.3 | cm |
| length range | [4, 10] | cm |
| initial fly distance from trench | [0, 2] | cm |

**Table 15:** Architecture of the high-level controller network for the vision-guided flight task. Visual input is pre-processed by a convolutional visual module and concatenated with the rest of the observations. Layer sizes indicated in parentheses. This high-level controller outputs a steering command of dimension 7 × (future steps + 1). The steering command is then concatenated with the proprioception and vestibular (but not task-input and visual) components of the observations and sent as input to the low-level controller network, which is the policy from the flight imitation task (see Table 11).

| *input: observations* |  |
| --- | --- |
| Linear (256) |  |
| LayerNorm |  |
| tanh |  |
| Linear (256) |  |
| ELU |  |
| Linear (128) |  |
| ELU |  |
| Linear (42) | Linear (42) |
|  | softplus |
| *output: action mean* | *output: action std* |

### Modeling walking behavior

#### Reference walking data preparation

We obtained single-camera top-down view videos of multiple freely behaving *Drosophila* with 2D keypoint tracking from Robie et al., (Extended Data Fig. 6A). Briefly, groups of 10 walking flies were recorded at 150 fps in a shallow, flat-bottomed, 50 mm diameter arena. The 2D positions of 17 keypoints were predicated with the Animal Part Tracker (APT) ^3^. From 9 such videos, we prepared the walking reference dataset. We used 13 of the 17 keypoints: three on the head, three on the thorax, one at the tip of abdomen and the 6 leg tips, as shown in Extended Data Fig. 6. We selected female flies and isolated walking trajectory segments based on the following criteria. At each frame, we required: (i) the distance to the other flies in the arena is larger than one body length, (ii) the velocity component parallel to the fly body is larger than perpendicular to the body. Then we required (iii) snippet duration of at least 20 frames (133 ms, roughly corresponds to one fly step), and (iv) ratio of mean leg tip speed to mean center-of-mass speed smaller than 1.5. This produced a set of trajectory snippets with flies walking at different speeds, turning, and standing still. The average walking speeds per snippet are distributed approximately in the range [0, 4] cm/s (Fig. 3e, inset). We then linearly interpolated the walking snippets from 6.7 ms timesteps to the walking simulation control of 2 ms timesteps.

**Extended Data Figure 6:**
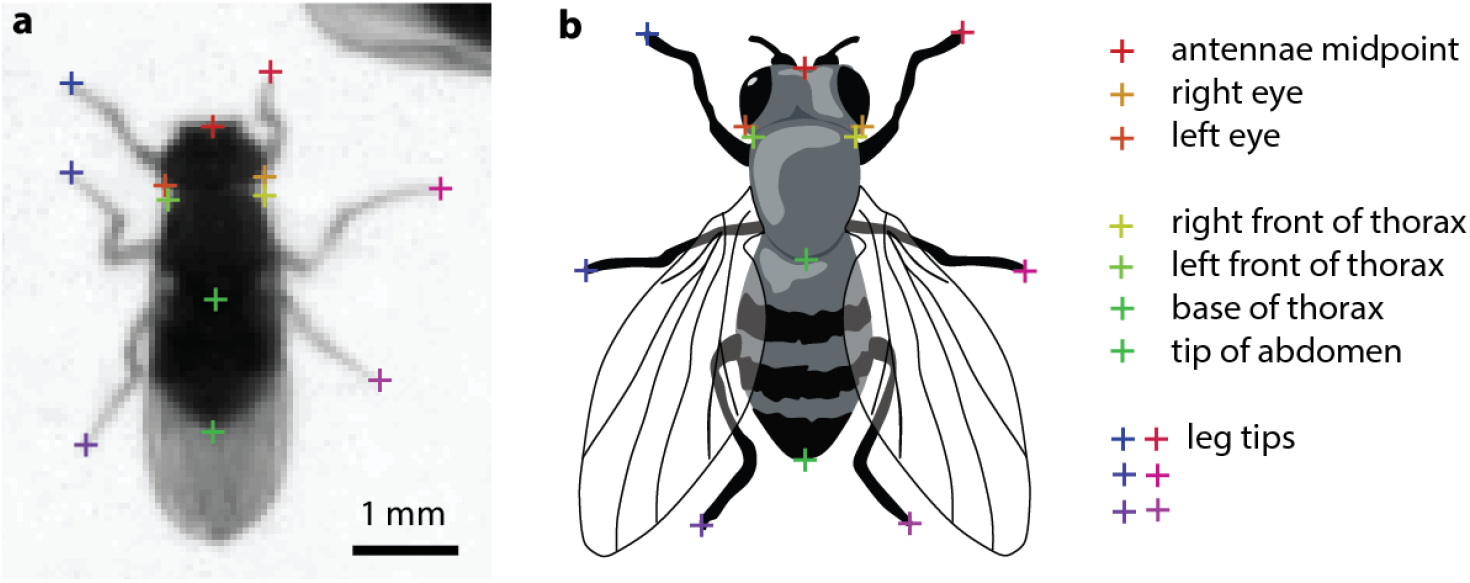
2D keypoints used to track fly walking. (**a**) A top-down view video frame of a walking fly with keypoints inferred with Animal Part Tracker (APT). (**b**) Keypoint definition.

The 13 keypoints tracked in two dimensions, however, are not sufficient for the RL task reward calculation (described below), which requires complete specification of the model’s pose, position, and orientation. Specifically, this full-body represention should include all joint angles, joint axis orientations, joint velocities, body position and orientation, and leg tip positions. Obtaining the full-body data for the model from the experimental data required, first, to lift the 2D walking snippets by complementing the horizontal *x, y*-coordinates with the third vertical *z*-dimension. Based on a separate side-view video of a walking fly [Akitake et al., 2015] and on our fly model’s default standing position, we approximated the body height and pitch angle during walking by a single fixed value. From this video, we also estimated the amplitudes of the arcs traversed by leg tips during swing motion. The amplitudes were *A* = 0.086, 0.047, 0.051 cm for the T1, T2, T3 legs, respectively. We approximated the *z*-coordinate of the leg-tip swing arcs by the sine function as *z* = *A* sin(*x*), with *x* going from 0 to *π* for each single leg swing. Using the 2D coordinates of the leg tip keypoints, we separated leg swings from stances based on the leg tip horizontal velocities in fly’s egocentric reference frame. Then we added the approximate sine-arcs to the swing segments of the leg tip trajectories, while we kept *z* = 0 for the stances. This procedure produced 3D coordinates for the 13 keypoints in the walking snippets selected earlier.

As a final step, we computed the full-body reference poses for each frame in all the snippets. We added to the fly model equivalent 13 keypoint sites and performed inverse kinematics fitting of the whole model body to the fly poses in the 3D walking snippets. For each snippet, we rescaled the reference keypoints to match the size of the fly model. In each frame, we fit simultaneously all 13 keypoints by minimizing the following objective with respect to the model joint angles, **q** = (*q*_1_, *q*_2_, …):

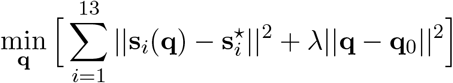

where **s**_*i*_(**q**) and 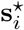 are the 3D Cartesian coordinates of the 13 keypoints of the model and the fitting target pose, respectively. We used gradient descent to minimize this objective. To utilize the time-continuity across frames, we used the final pose fitted for the previous frame to initialize the fitting procedure for each subsequent frame. As the 13 keypoints (only 6 of which specify the leg tips) do not fully define the leg postures in space, we added a small regularization term to encourage fitting poses that are closer to the default standing pose of the fly model. The default pose is specified by a vector of model joint angles, **q**_0_, a vector of zeros in our case. The regularization strength is *λ* = 1× 10^−4^ cm^2^*/*rad^2^. Having found the joint angles **q** for the reference poses in each frame, we also computed joint velocities, *d***q***/dt*, using finite differences.

This procedure resulted in a complete full-body representation (joint angles, positions, orientations, velocities) of the reference walking trajectories. This is our reference data for the walking imitation task. In total, the walking dataset is comprised of ∼ 16,000 walking snippets, amounting to ∼ 80 minutes of fly walking behavior. The dataset is available at Supp. Data.

#### Walking imitation task configuration

The reinforcement learning task is set as follows. In each episode, a reference trajectory is randomly selected from the walking dataset described above. The model is required to track the CoM position and orientation of the reference fly body, as well as the detailed motion of the legs. The initial model posture and the CoM position and velocity are set equal to the reference in the first step of the selected trajectory. The episode ends when the end of the trajectory is successfully reached or terminates early if the model CoM is displaced from the reference by more than 0.3 cm (one body length.)

We used a multiplicative version of the imitation reward mostly similar to [Peng et al., 2018, Merel et al., 2020b]. At every timestep, the reward measures the similarity between the current model and reference pose, position, and orientation. The reward function is formulated as a product of (not normalized) Gaussian functions, one Gaussian for each quantity compared, e.g. a joint angle, the CoM position, etc. As the Gaussians are not normalized, each of them is in the range [0, 1], and so is the total multiplicative reward. There is no unique way to construct this imitation reward function and some of the terms could be redundant/overlapping. At each time step, the reward *R* is calculated as:

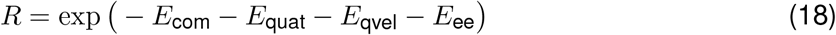

where the four terms expressing different aspects of the pose tracking objective are:

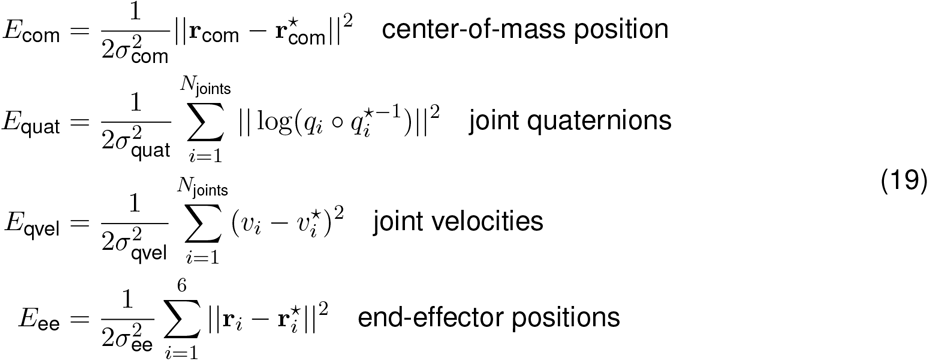

*E*_com_ measures the similarity between the current model and reference Cartesian CoM positions, **r**_com_ and 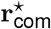. *E*_quat_ measures the similarity between the model and reference joint quaternions *q*_*i*_ and 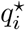 for all joints, including the root joint. Joint quaternions encode both the joint angle and the direction of the joint rotation axis for hinge joints (or body orientation for the root joint.) *E*_qvel_ expresses the similarity between the model and reference joint velocities, *v*_*i*_ and 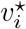 compares the Cartesian positions of the end-effectors (six leg tips), **r**_*i*_ and 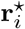.Each one of the four error *E* terms has its own reward tightness (scale) hyperparameter, expressed as a standard deviation *σ*. See Table 18 for the hyperparameter values used.

In the walking imitation task, we set the wings in the retracted position. We removed the actuators of the wings, proboscis, antennae. We retained their DoFs in the model but excluded these DoFs from the policy observations. We didn’t use vision in this task. The observation and action dimensions were 741 and 59, respectively, see Table 16 and Table 17 for detail. The policy network architecture is shown in Table 19. We attached MuJoCo adhesion actuators to leg tips to simulate fly’s adhesive pads. The control semantics for the adhesion actuators is the required adhesion force, ranging between zero to one fly body weight for each leg. The rest of the actuators were position actuators receiving target joint angles as control from the policy. We didn’t use any data related to adhesion during walking and didn’t constrain the model as to how to use the adhesion actuators (e.g., there are no adhesion terms in the reward.) We observed, however, that the agent generally preferred to activate the adhesion while the legs were in stance to increase friction with the ground. In this task, we did not use the DMPO action penalization because the full-body imitation reward automatically constrains the policy outputs (target joint angles) to the proper ranges.

**Table 16:** Observations in the walking imitation task. All observables are calculated in fly’s egocentric reference frame. The actuator activation state units are the same as the corresponding actuator control units.

| Category | Observables (policy inputs) | Units | Array shape |
| --- | --- | --- | --- |
| vestibular | velocimeter | cm/s | 3 |
|  | accelerometer | cm/s<sup>2</sup> | 3 |
|  | gyro (rotational velocity) | rad/s | 3 |
|  | gravity direction | cm | 3 |
| proprioception | joint angles | rad | 85 |
|  | joint velocities | rad/s | 85 |
|  | actuator activation state | same as control | 59 |
|  | end-effector positions | cm | (7, 3) |
| mechanoreception | leg force sensors | dyn | (6, 3) |
|  | leg touch sensors | dyn | 6 |
| task input<br>(steering commands) | reference CoM displacement | cm | (65, 3) |
|  | reference orientation displacement | unitless quaternion | (65, 4) |
| total obs. dimension |  |  | 741 |

**Table 17:** Actions in the walking imitation task.

| Actions (policy outputs) | Array shape |
| --- | --- |
| head | 3 |
| abdomen | 2 |
| legs | (6, 8) |
| leg adhesion | 6 |
| total action dimension | 59 |

**Table 18:** Walking imitation task hyperparameters.

| Parameter | Value | Units |
| --- | --- | --- |
| physics_timestep | $2 \times 10^{-4}$ | s |
| control_timestep | $2 \times 10^{-3}$ | s |
| joint_filter | 0.01 | s |
| adhesion_filter | 0.007 | s |
| claw_friction | 1.0 |  |
| future_steps | 64 |  |
| time_limit | 10.0 | s |
| terminal_com_dist | 0.3 | cm |
| reward parameters: |  |  |
| $\sigma_{com}$ | 0.0785 | cm |
| $\sigma_{qvel}$ | 53.780 | rad/s |
| $\sigma_{ee}$ | 0.0735 | cm |
| $\sigma_{quat}$ | 1.225 | rad |

**Table 19:** Walking policy network architecture. Layer sizes indicated in parentheses.

| *input: observations* |  |
| --- | --- |
| Linear (512) |  |
| LayerNorm |  |
| tanh |  |
| Linear (512) |  |
| ELU |  |
| Linear (512) |  |
| ELU |  |
| Linear (512) |  |
| ELU |  |
| Linear (59) | Linear (59) |
|  | softplus |
| *output: action mean* | *output: action std* |

As in the flight imitation task, in addition to the standard set of egocentric vestibular and proprioception observables, the policy receives task-specific inputs: the Cartesian CoM displacement and the orientation (quaternion) displacement of the reference trajectory with respect to the model at the current timestep plus 64 timesteps into the future. In contrast to the flight tasks, the actions output by the policy are not fed to the actuators directly but first undergo filtering (averaging) in time with time constants `joint_filter` and `adhesion_filter` for joint and adhesion actuators, respectively (see Table 18). As a result, the current actuator target joint angles (or target force in adhesion actuators) are generally different from the current policy control output and therefore considered as the actuator’s internal state. This internal state is added as an additional observable in this task, see Table 16. As in the flight imitation task, the trained walking policy can be used as a low-level walking controller with the two task-specific inputs (CoM and quaternion displacements) serving as high-level steering control commands.

### Adhesion, friction, and contact forces during walking

The fly model’s ability to attach to and to walk on inclined surfaces is enabled by the combination of adhesion, friction, and contact forces. In this section we describe the details of the adhesion mechanism in MuJoCo and how the fly model uses the leg adhesion actuators. Let’s consider a simple example of the stationary fly on an inclined plane. When a contact is detected between a tarsal claw collision geom and the floor surface, MuJoCo computes the (constraint) contact force, or the ground reaction force in this case (Extended Data Fig. 7a). Within the Coulomb friction model, as long as the contact force vector **f**_contact_ is within the friction cone boundaries, the tangential component of the net external force (**f**_weight_, the fraction of the total fly weight supported by the given leg in this simplified example), which acts to produce slipping motion, will be balanced by the tangential component 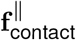 of the contact force. The (elliptic) friction cone^4^ includes all contact force vectors satisfying 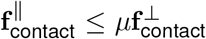 where *µ* is the static friction coefficient. Outside of the friction cone, i.e. for contact forces beyond the threshold 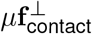,slipping motion will occur. Note that within this friction model, the cone angle is a function of the friction coefficient alone and is given by *θ* = tan^−1^ *µ*. In our model, *µ* = 1 and *θ* = 45^°^.

In MuJoCo, the action of an adhesion actuator is equivalent to injecting force in the normal contact direction (Extended Data Fig. 7a) effectively acting to push the fly’s claw into the floor. In response, the normal component 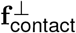 of the contact force will increase by the same amount. While there is no change in the tangential contact force component due to the adhesion, note how the net contact force vector is now further away from the friction cone boundary thus providing a larger slip resisting margin which can be used for, e.g., forward walking propulsion. Beyond its role in the Coulomb friction mechanism, the adhesion force can also directly counteract gravity to enable walking on arbitrarily oriented surfaces, such as vertical walls or ceiling.

Extended Data Fig. 7b shows the time-lapse of the walking task where the fly model was trained to use adhesion to overcome bumpy terrain. This task is similar to the walking imitation task described above. The fly is required to imitate a single real-data walking snippet of walking straight at fixed speed 2.7 cm/s. In fly’s way, we introduced a sine-like obstacle that cannot be overcome without adhesion. The bump obstacle is procedurally regenerated at each training episode with the bump’s height and length varying in the ranges [0, 2] and [2, 4] cm, respectively. Thus the bump inclination angle was between 0^°^ – 72^°^. We also added a small action penalty, `epsilon_penalty` = 3 ×10^−4^, through the DMPO agent mechanism, to encourage the agent to prefer economic actions, including the adhesion action. We recorded the adhesion action and contact forces during a trained policy rollout on a bump with max inclination angle of ∼ 45°, as shown in Extended Data Fig. 7(c-f).

The adhesion forces produced by the leg adhesion actuators while overcoming the obstacle are shown in Data Fig. 7c. The model’s use of adhesion increases as the terrain angle becomes steeper. It is evident that the fly model learned to use mostly the T1 and T2 leg pairs on the way uphill, and mostly T3 on the way downhill. Due to the lack of constraints, there is an asymmetry (degeneracy) between the left and right leg adhesion utilization, which we didn’t attempt to resolve. In out model, the largest adhesion force per leg is one fly body weight, which is also shown in the figure for comparison. The norm of the corresponding contact force vectors |**f**_contact_| for each leg is shown in Extended Data Fig. 7d. The effect of creating a larger slip-resisting margin – moving **f**_contact_ further away from the friction cone boundary – with increasing adhesion is shown in Extended Data Fig. 7e. Without the adhesion, most of the leg-floor contacts would not have been able to counteract the slipping force load, especially in the “driving” T1, T2 legs on the way up, and T3 on the way down (Extended Data Fig. 7f).

**Extended Data Figure 7:**
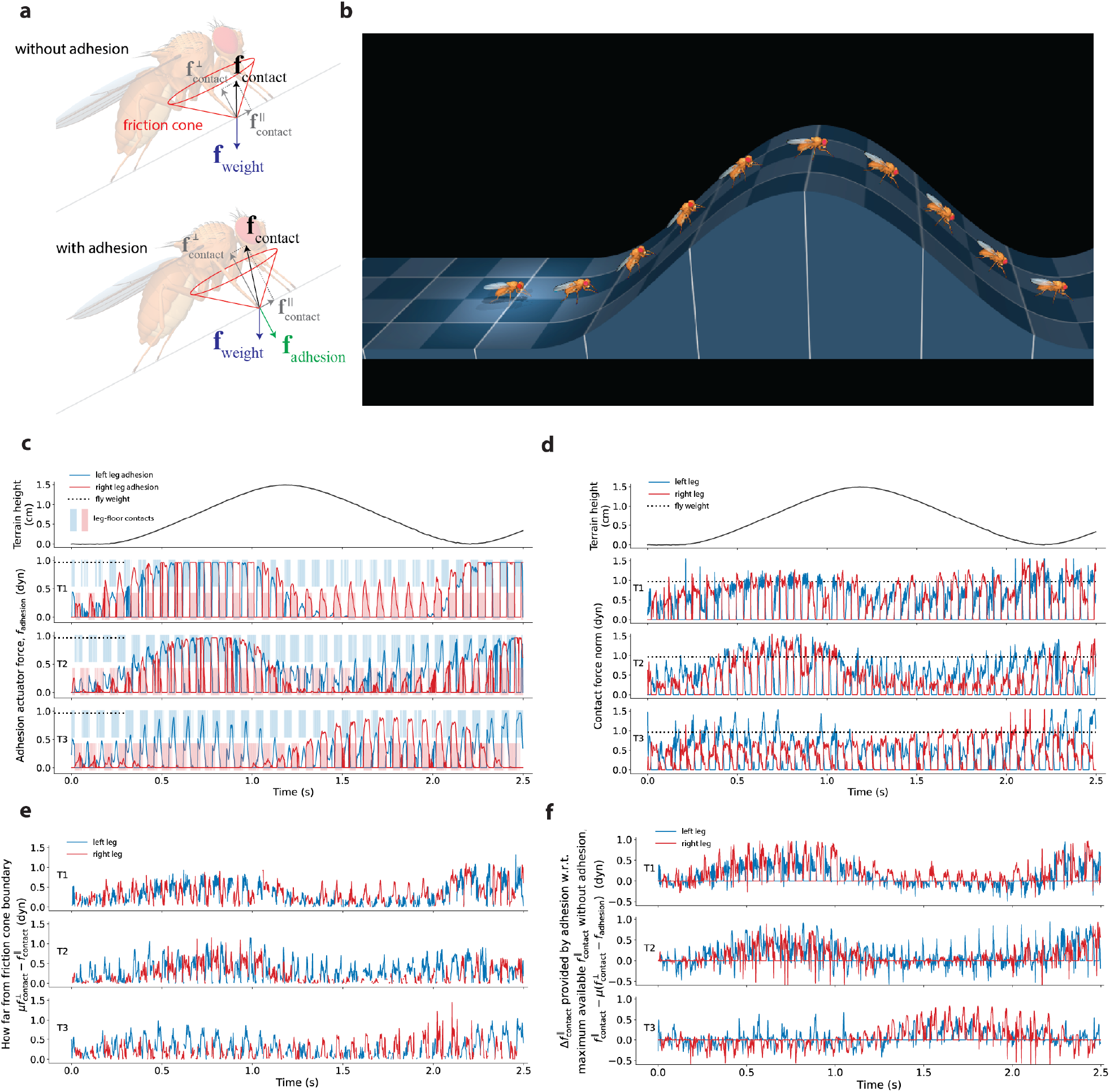
Adhesion and contact forces during walking on hilly terrain. (**a**) Schematic diagram of leg-floor contact forces for the fly model stationary standing on an inclined surface. The adhesion actuator injects force in the normal direction, which in response increases the normal component of the contact force. This creates a larger margin between the tangential contact force component which resists slipping and the slip threshold (the friction cone boundary). (**b**) Time-lapse of the trained policy rollout of the RL task where the fly model learns to use the adhesion mechanism to overcome sine-like hills. All the following panels correspond to this policy rollout. (**c**) Adhesion actuator forces generated by the fly’s claws during the policy rollout shown in **b**. In our model, the largest adhesion force per leg is one fly body weight. The fly body weight, *mg* = 0.97 gr, is shown for comparison. Also the leg-floor contacts are shown for clarity. (**d**) Contact force norm during the policy rollout. The fly weight is shown for comparison. (**e**) The difference between the slip threshold force and the tangential component of the contact force. This is the “margin” available to resist slipping under external forces and propulsion generation.(**f**) The difference between the actual tangential contact force and the largest tangential contact force that would have been available without adhesion. Positive means the contact would have slipped without adhesion. Negative means the contact would not have slipped without adhesion (that is, the contact is inside the friction cone already without adhesion).

## Acknowledgments

This work was supported by the Howard Hughes Medical Institute and Google DeepMind. The authors thank Piotr Trochim for writing the Blender-to-MuJoCo export plug-in, Ben Moran for open-sourcing the plug-in, Nimrod Gileadi for reviewing code submissions to `dm_control`. Thomson Rymer, Emily Tenshaw (Janelia Project Technical Resources) and Tyler Paterson (Janelia Project Pipeline Support) for performing APT annotations. Janne Lappalainen, Diptodip Deb, and the members of the John Tuthill lab and the Bing Brunton lab for critical reading of the manuscript and stimulating discussions. ZS was supported by the German Research Foundation (DFG) through SPP 2041, Germany’s Excellence Strategy (EXC-Number 2064/1, Project number 390727645) and the German Federal Ministry of Education and Research (BMBF; Tübingen AI Center, FKZ: 01IS18039A). ZS is a member of the International Max Planck Research School for Intelligent Systems (IMPRS-IS).

## Declarations

The authors declare no competing interests.

## Supplement

### A Constructing the MuJoCo physics model

Here we describe the steps taken in order to create the physically-simulatable MuJoCo fly model, given the geometrical Blender model and measured masses. The resulting MuJoCo model is available at https://github.com/TuragaLab/flybody.

#### Initial conversion

The Blender-to-MuJoCo export plug-in, available at https://github.com/google-deepmind/dm control, was used to export a raw MuJoCo model containing only geometrical information: body meshes and a kinematic tree with joint axes and limits.

#### Model building script

The raw MuJoCo model was then loaded and manipulated with a Python script using PyMJCF, a Python library for model manipulation which is part of Google DeepMind’s dm control suite [Tunyasuvunakool et al., 2020]. The following steps were taken.

1. Enforced consistent naming everywhere using a `part sternum side` convention e.g., “ `coxa T3 right`”. Consistent naming allows for conveniently readable loops like 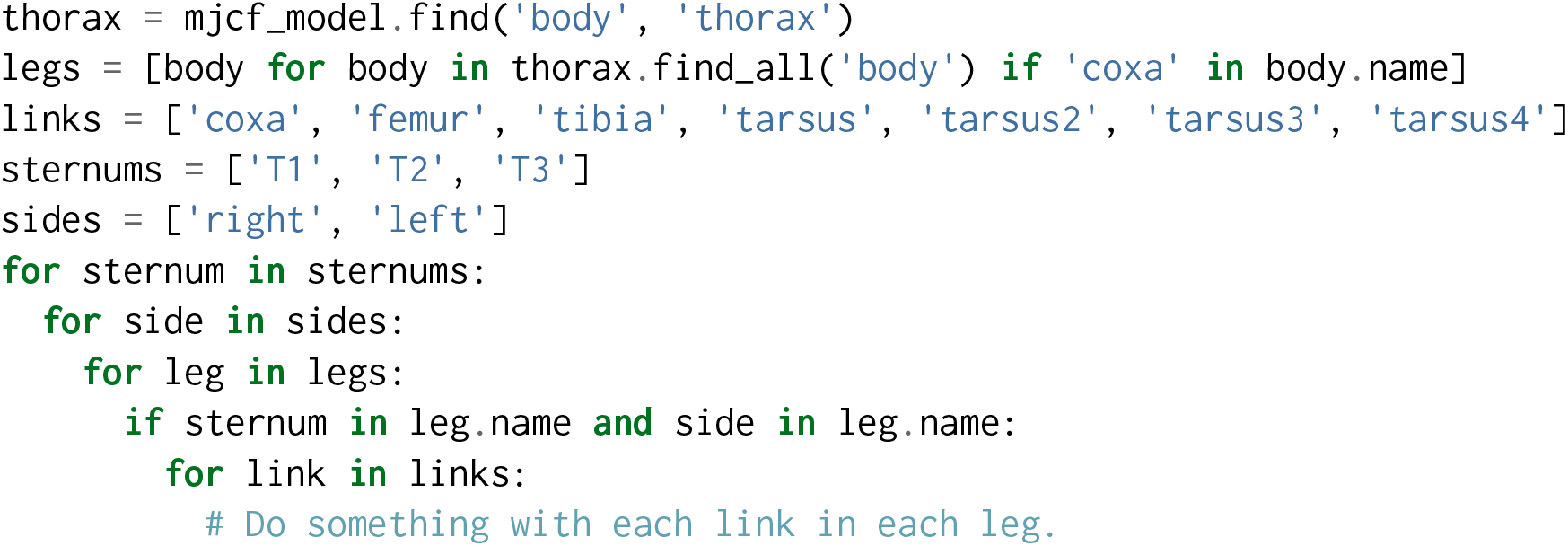
2. Made use of MuJoCo’s cascading defaults mechanism, avoiding repeated values. Common properties like angle ranges, damping and actuator properties for all joints of the same type or masses of links of the same type are given by a single number in the model, defined in a default class, which is then inherited by elements in the kinematic tree.
3. Model units were chosen to be CGS, for better numerical precision. Note that in MKS, some values are extremely small, for example the inertia of a tarsus link is on the order of 1 ×10^−20^ kg m^2^, while in CGS is 1 ×10^−13^ g cm^2^. The difference in accuracy is significant for double-precision floating-point arithmetic.
4. Body moments-of-inertia were computed by MuJoCo given mesh geometries and empirical masses, assuming uniform density for each body part.
5. Kinematic symmetry was enforced to numerical precision, transverse of the sagittal plane.
6. Joint axis orientations were reflected through the saggital plane, ensuring identical semantics on both sides i.e., joint rotation in the positive (negative) direction always corresponds to extension (flexion) and abduction (adduction), respectively.
7. Joint angle reference values were chosen so that 0.0 corresponds to the base pose, see Fig. 1.
8. A layer of primitive collision geoms was created, initially by letting MuJoCo fit primitives to meshes by matching ineritas, and then by manual fine-tuning.
9. Manually excluded pairs of bodies that cannot collide, both to avoid spurious collisions and to increase simulation speed. Note that certain collisions that are possible in real flies were also excluded. In particular since the modeled wings are rigid and while real wings are flexible, wing-wing and wing-body collisions cannot be well modeled.
10. Added tendons to the abdomen and the tarsi. These “fixed” tendons are a simplification of spatially routed tendons, whose length corresponds to a linear combinations of joint angles, allowing a single actuator to act on multiple joints e.g., to flex the multiple links of the tarsus or abduct the entire abdomen. See examples in Fig. 1.
11. Set the default body pitch angle to 47.5°, following [Muijres et al., 2014]. Re-orient the wing joint axes such that when the body is at 47.5°, the wing yaw axes are strictly vertical and the stroke plane is horizontal.
12. Added a total of 78 actuators:
  - 8 actuators in each leg (coxa: 3, femur: 2, tibia: 1, tarsus: 2), using desired angle (position) semantics, with gains chosen so that a force of approximately one body weight can be applied at the end-effector at the base pose.
  - Wings actuators (yaw, roll, pitch) have torque semantics with gain values of 18.0 dyn cm, see Methods.
  - 16 additional actuators for proximal joints: head and rostrum (4), haustelli (2), labri (2), antennae (6) and abdomen (2).
  - 6 adhesion actuators at the claws which can apply a force up to 1× body weight.
  - 2 adhesion actuators in the mouth (labrum).
13. Added egocentric sensors (also see Table 20):
  - An ideal accelerometer and gyro to the thorax, corresponding to processed vestibular sensor information.
  - An ideal velocimeter, corresponding to processed information from air motion sensors in the hair follicles.
  - Added force and touch sensors at the end effectors. The former report the force passing through the first tarsus joints, while the latter report pressure applied to the claw.
  - Two eye cameras, at the geometric center of the eyes. These are standard openGL cameras with a wide 140°field-of-view, see main text for how these were used in visually guided experiments.

### B Sensory system components

**Table 20:** Correspondence between the sensory system components in *Drosophila* and in our fly model.

| **Drosophila sense organ, receptor** | **Sensory modality** | **Purpose, detection, sensing** | **Corresponding fly model sensor** | **References** |
| --- | --- | --- | --- | --- |
| Compound eyes | Light, photoreception | Self-movement (optic flow, optomotor response), looming stimuli | Eye cameras, velocimeter, gravity direction | [Hengstenberg, 1993, Strausfeld and Seyan, 1985] |
| Ocelli | Light, photoreception | Dorsal light response, body orientation, postural correction | ditto | [Hengstenberg, 1993, Strausfeld and Seyan, 1985, Mimura et al., 1970] |
| Halteres (modified 2nd wing pair) | mechanoreception | Coriolis forces during flight; balance/equilibrium organ | Gyro, accelerometer | [Strausfeld and Seyan, 1985, Fayyazuddin and Dickinson, 1996] |
| Leg campaniform sensilla | mechanoreception | Load/gravity | Force, touch, gravity direction | [Dinges et al., 2021] |
| Johnston's organ (2nd antennal segment) | Mechanoreception (exteroception) | Gravity, air flow, sound (courtship song) | Gravity direction, velocimeter, sound not implemented | [Kamikouchi et al., 2009, Mimura et al., 1970, Mamiya and Dickinson, 2015] |
| Neck proprioceptors (prosternal organ, CO) | Mechanoreception (proprioception) | head position; postural correction | Joint angles, joint velocities | [Strausfeld and Seyan, 1985] |
| Wing base tegula | Mechanoreception via a hair plate and campaniform sensilla | Wing position (proprioception) | ditto | [Fudalewicz-Niemczyk, 1963] |
| Femoral chordotonal organ (FeCO) | mechanoreception | Flexion, extension of the tibia (proprioception); vibration (exteroception) | ditto | [Mamiya et al., 2023] |
| Coxa and trochanter hair plates | mechanoreception | Registering extreme positions of the legs (proprioception) | ditto | [Murphey et al., 1989, Merritt and Murphey, 1992] |
| Wing base campaniform sensilla | mechanoreception | Wing twist, wing load | *Not implemented* | [Cole and Palka, 1982, Dickinson and Palka, 1987] |
| Wing veins campaniform sensilla | mechanoreception | Aeroelastic deformations of wing | *Not implemented* | [Cole and Palka, 1982, Dickinson and Palka, 1987] |
| Wing margin chemosensory bristles | chemoreception | Airborne molecules (odor plumes, pheromones) | *Not implemented* | [Houot et al., 2017] |
| Arista (4th-6th antennal segments) | Hygro-, chemo-, thermoreception | Odor, pheromone plumes, temperature gradients | *Not implemented* | [Foelix et al., 1989] |
| Labrum and tarsal gustatory receptors | Hygro- and chemoreception | Chemical compounds in the substrate | *Not implemented* | [Montell, 2009] |

### C Simulation speed

The timing of different components within one control timestep of the simulation is shown in Table 21. We simulated the different fly model behaviors on a single core of Intel Xeon CPU E5-2697 v3 @ 2.60GHz. The total simulation control step time is broken down into its components as *Total step time* = *Policy* + *RL env* + *MuJoCo control*, where *Policy* is the forward pass through the policy network, *RL env* is the dm control python RL environment, and *MuJoCo control* is the total of MuJoCo physics timesteps per one RL environment control timestep [Tunyasuvunakool et al., 2020]. The policy forward pass was run on CPU, same as in actors during training. % real time is shown for both the total simulation timestep and for the MuJoCo component alone. The averages are calculated from 10,000 simulation steps with std-to-mean ratios being on the order of ∼ 0.01 For vision-guided flight, “bumps” task times are shown; “bumps” and “trench” tasks perform similarly; visual input was rendered on an NVIDIA Titan Xp GPU.

**Table 21:**
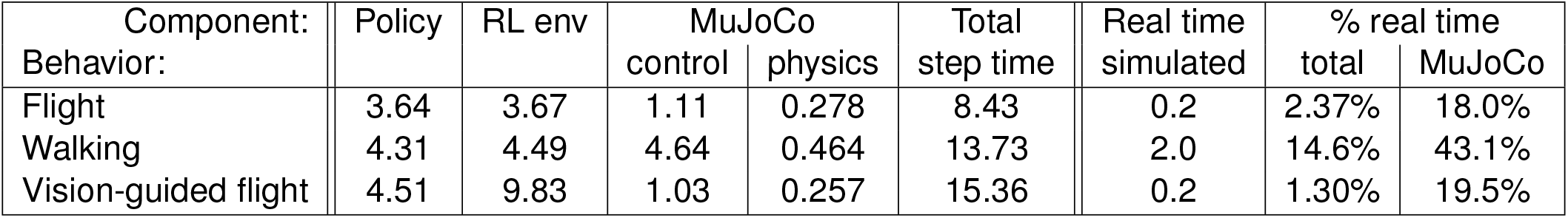
Break down of average times of one control timestep for different fly model behaviors simulated on a single core of Intel Xeon CPU E5-2697 v3 @ 2.60GHz. All times are in units of ms.

| Component:<br>Behavior: | Policy | RL env | MuJoCo |  | Total<br>step time | Real time<br>simulated | % real time |  |
| --- | --- | --- | --- | --- | --- | --- | --- | --- |
|  |  |  | control | physics |  |  | total | MuJoCo |
| Flight | 3.64 | 3.67 | 1.11 | 0.278 | 8.43 | 0.2 | 2.37% | 18.0% |
| Walking | 4.31 | 4.49 | 4.64 | 0.464 | 13.73 | 2.0 | 14.6% | 43.1% |
| Vision-guided flight | 4.51 | 9.83 | 1.03 | 0.257 | 15.36 | 0.2 | 1.30% | 19.5% |

In addition, note that the simulation speed can be improved further:

- When not flying, a much larger time-step can be used.
- When flying, contacts can (sometimes) be ignored.
- MuJoCo simulations can be easily duplicated across threads.
- As of MuJoCo 3.0, simulation is supported on GPU and TPU accelerators which can achieve much higher throughput, especially for RL training, as both the physics and network are on the same processor. We have not yet attempted to run our environments on accelerators as this would require rewriting the environment logic (standard Python is bound to CPU).

## Footnotes

1 https://mujoco.readthedocs.io/

2 https://github.com/willdickson/fmech

3 https://github.com/kristinbranson/APT

4 Our fly model uses elliptic friction cones. MuJoCo also supports pyramidal friction cones.

